# Distinct structural and functional heterochromatin partitioning of lamin B1 and B2 revealed using genome-wide Nicking Enzyme Epitope targeted DNA sequencing

**DOI:** 10.1101/2024.07.11.603107

**Authors:** Sagnik Sen, Pierre-Olivier Estève, Karthikeyan Raman, Julie Beaulieu, Hang Gyeong Chin, George R. Feehery, Udayakumar S. Vishnu, Shuang-yong Xu, James C. Samuelson, Sriharsa Pradhan

## Abstract

A genome-wide chromatin profiling technology, named as Nicking Enzyme Epitope targeted DNA sequencing (NEED-seq) in which antibody-targeted controlled nicking by Nt.CviPII-pGL is used to study specific protein-DNA complexes. NEED-seq is performed *in situ* in formaldehyde fixed cells, allowing for both visual and genomic resolution of epitope bound chromatin. When applied to nuclei, NEED-seq yielded genome-wide chromatin associated proteins and histone post-translational modifications (PTMs). NEED-seq of lamin B1/B2 demonstrated their association with heterochromatin. Lamin B1 and B2 associated domains (LAD) segregated to three different states, and states with stronger LAD correlated with heterochromatic marks. Hi-C analysis displayed A and B compartment with equal lamin B1/B2 distribution, although methylated DNA remained high in B compartment. LAD clustering with Hi-C resulted in subcompartments, with lamin B1-B2 partitioning to facultative and constitutive heterochromatin respectively and were associated with neuronal development. Thus, lamin B1 and B2 have structural and functional partitioning in mammalian nucleus.

**Figure 1:**
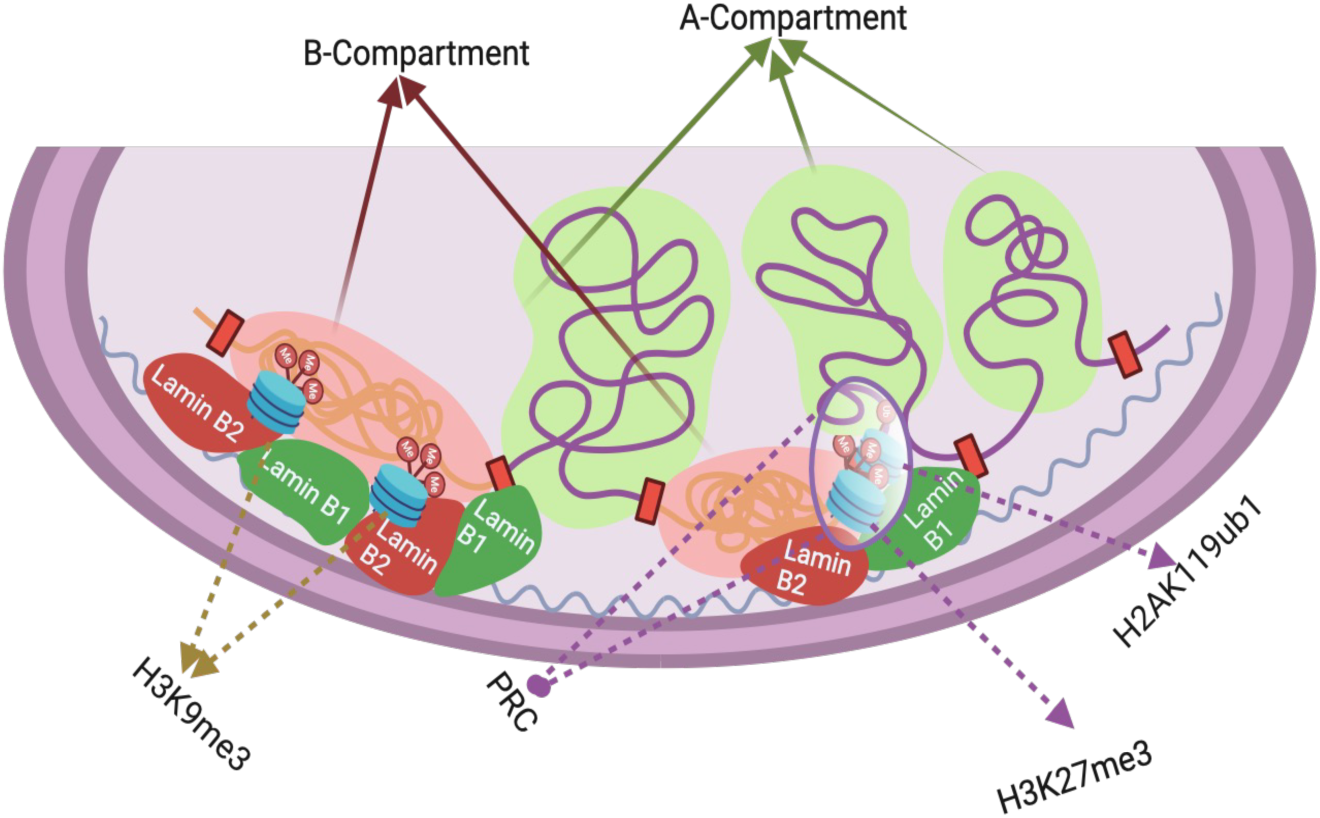
**Graphical abstract:** Model depicting association of lamin B1 and B2 in A (facultative heterochromatin) and B (constitutive heterochromatin) compartmentalization.

## Introduction

Epigenetic inheritance is documented during physiological processes including cell division, and mammalian development (Probst et al, 2009; Kim et al., 2009; Xu and Xie, 2018). Aberrant epigenetic patterns lead to pathogenesis of various human diseases, including cancer (Berdasco and Esteller, 2010; Zoghbi and Beaudet, 2016). In combination with next-generation sequencing (NGS) technologies, genome-wide epigenetic feature profiling has enabled an unprecedented opportunity to decipher the gene expression profile in normal and diseased cells (Sarda and Hannenhalli, 2014; Cieślik and Chinnaiyan, 2018; Nones and Patch, 2020). Primarily, a combination of the transcription factors (TFs) occupancy on the DNA, post-translational modifications (PTMs) of histones, and DNA cytosine methylation pattern determines the gene expression output (Partridge et al., 2020; Bowman and Poirier, 2015; Bell et al., 2011; Anastasiadi et al., 2018). Therefore, genome-wide TF occupancy, histone, and non-histone post-translational modification mapping have become a central focus of biological studies. For these studies, there are both antibody independent and dependent methodologies. The antibody-independent method, for example DamID, involves creating a functional fusion protein by tethering a DNA-binding protein to *E. coli* Dam methyltransferase (Dam). Once expressed in cells, methylation of adenine occurs in GATC sequence of DNA near the native binding sites of the Dam fusion partner. Since adenine methylation does not occur endogenously in most eukaryotes, it provides a unique tag to mark protein interaction sites (van Steensel et al., 2001). These sites can be mapped genome-wide using an N6-adenine methyl-specific restriction enzyme DpnI. This technology has limited resolution due to its dependency on GATC sites in the vicinity of the DNA binding protein. This method has been applied successfully in Drosophila using a Pol II-Dam fusion (Southall et al., 2013). In another antibody-independent method, the ChEC method (chromatin endogenous cleavage), involves expressing fusion proteins in cells, where micrococcal nuclease (MNase) is C-terminally fused to the DNA binding proteins of interest. The bifunctional nucleases are activated *in situ* with Ca^2+^ cations to locally introduce double-stranded DNA cleavage (Schmid et al., 2004). ChEC-seq maps protein binding with 100-200 bp resolution with high specificity.

In the antibody-dependent method, the most utilized technology is chromatin immunoprecipitation followed by DNA sequencing (ChIP-seq, Johnson et al., 2007). This technique could be adapted to a wide array of transcription factors, polymerases and transcriptional machinery, structural proteins, protein modifications, and DNA modifications, with specific antibodies (Park, 2009). There are also various improvements of ChIP-seq to accommodate low cell numbers, including single cell (Grosselin et al., 2019). ChIP-seq relies on formaldehyde crosslinked cells, chromatin fragmentation and solubilization as the starting immunoprecipitation material. Typically, an antibody of interest is added to the lysate, and the antibody-bound chromatin is captured. The DNA is purified for library preparation and sequencing, allowing base-pair resolution mapping of TFs (Rhee and Pugh, 2011; Skene and Henikoff, 2015). Another variation of ChIP without formaldehyde cross-linking, known as native-ChIP, is also used (Cosseau et al., 2009). However, both technologies have advantages and disadvantages. Although native-ChIP minimizes problems with epitope masking, it can suffer from incomplete extraction efficiency of protein DNA complexes and the potential loss of weakly interacting proteins on DNA.

Antibody based modified ChIC-seq (chromatin immunocleavage) was developed to retain the advantages of enzyme tethering methods to specifically bound antibodies targeted to DNA binding proteins (Schmid et al., 2004). A key feature of this protocol is calcium-induced MNase cleavage on both sides of the epitope, the epitope-DNA complex is recovered for DNA extraction and sequencing. Another analogous technology known as cleavage under targets and release using nuclease (CUT&RUN), like the ChIC-seq technique, is versatile and utilizes non-cross-linked cells generally. This is simpler than native ChIP-seq while retaining the advantages of *in situ* application (Skene and Henikoff, 2017). CUT&RUN is shown to provide near base-pair resolution using ∼100 human cells for an abundant histone PTM and ∼1000 cells for a transcription factor (Skene et al., 2018). However, in a formaldehyde fixation study to freeze transient protein interactions in cells, CUT&RUN RNA Pol II signal at the TSS was ubiquitously reduced to ∼80% compared to the unfixed cells, without impacting the background signal (Miura and Chen, 2020). It was hypothesized that the formaldehyde fixation reduces the efficiency of chromatin cutting by pAG-MNase and/or affects the release of cleaved DNA fragments.

Here, we report a novel highly specific antibody dependent chromatin capture technology, Nicking Enzyme Epitope targeted DNA sequencing (NEED-seq), using a fusion nicking enzyme, Nt.CviPII-pGL (pGL, protein G plus linker domain), that retains the advantages of enzyme tethering methods, while providing both *in situ* visualization, and sequencing information of the target protein bound to chromatin generating functional genomics. NEED-seq has low background and high reproducibility. We negated non-specific DNA binding of the nicking enzyme fusion using heparin as a competitor. The epitope-bound nuclease is then activated by Mg^2+^ ions to locally introduce single-stranded DNA nicks (large DNA fragments were collapsed into small ones by frequent nicking). Nick-translation by DNA polymerase I in the presence of biotin and/or fluorophore conjugated dNTP reveals the precise visual and sequence location of the epitope. Mapping of such regions is shown to reveal proteins with a 100-200 bp resolution and excellent specificity. Using NEED-seq we have interrogated target histone PTMs, transcription factors, and structural proteins in a variety of cultured cells.

Structural proteins, lamin A and B are the major component of the nuclear lamina, interacting directly with DNA, linking the nuclear lamina to the cytoskeleton and chromatin. Lamin B1 and B2 are product of different genes and have high degree of sequence similarity. However, *Lmnb1* and *Lmnb2* knockout mice displayed neuronal migration in the developing brain defect (Kim et al., 2011). In a reciprocal knock-in mice experiment (higher production of lamin B1 from the *Lmnb2* locus and lower production of lamin B2 from the *Lmnb1* locus), the mice lines manifested neurodevelopmental abnormalities like those in conventional knockout mice, suggesting non-redundant, non-compensatory function of both proteins (Lee et al., 2014). In addition, lamin B1 and B2 regulate several cellular processes, such as tissue development, cell cycle, cellular proliferation, senescence, and DNA damage (Evangelisti et al., 2022). These experiments suggests that lamin B2 has specific role during development. To better understand how lamin B1 and B2 proteins differentially associate and regulate genomic regions, it is important to establish their relationship between epigenetic features and bound DNA. Here we have performed NEED-seq of lamin B1, B2, host of histone PTM, and genome-wide single base resolution DNA methylation analysis to determine the intra-and interconnectivity of lamin B1 and B2 with DNA and histone modification with special emphasis on heterochromatin compartmentalization.

## Results

### Nt.CviPII fusions

Nicking enzyme, Nt.CviPII (recognition sequence CCD, D = A, G and T), is an effective enzyme for genome wide accessible chromatin studies in formaldehyde fixed cells (Ponnaluri et al., 2017; Chin et al., 2020; Vishnu et al., 2021, Estève et al., 2020). We hypothesized that the nicking enzyme could be fused to antibody binding protein moiety, such as protein G, or L in combination or individually, for antibody directed DNA bound epitope visualization and mapping for both visual and functional epigenomics. We named this method as NEED-seq (Fig. 1). We fused protein G plus L at the C-terminus of Nt.CviPII and named the fusion as Nt.CviPII-pGL (Fig. S1A). The enzymes were purified to homogeneity (Fig. S1B, C). The fusion Nt.CviPII-pGL, displayed similar nicking activities on pUC19 plasmid as that of the Nt.CviPII enzyme alone (Fig. S2A, B).

**Figure 1:**
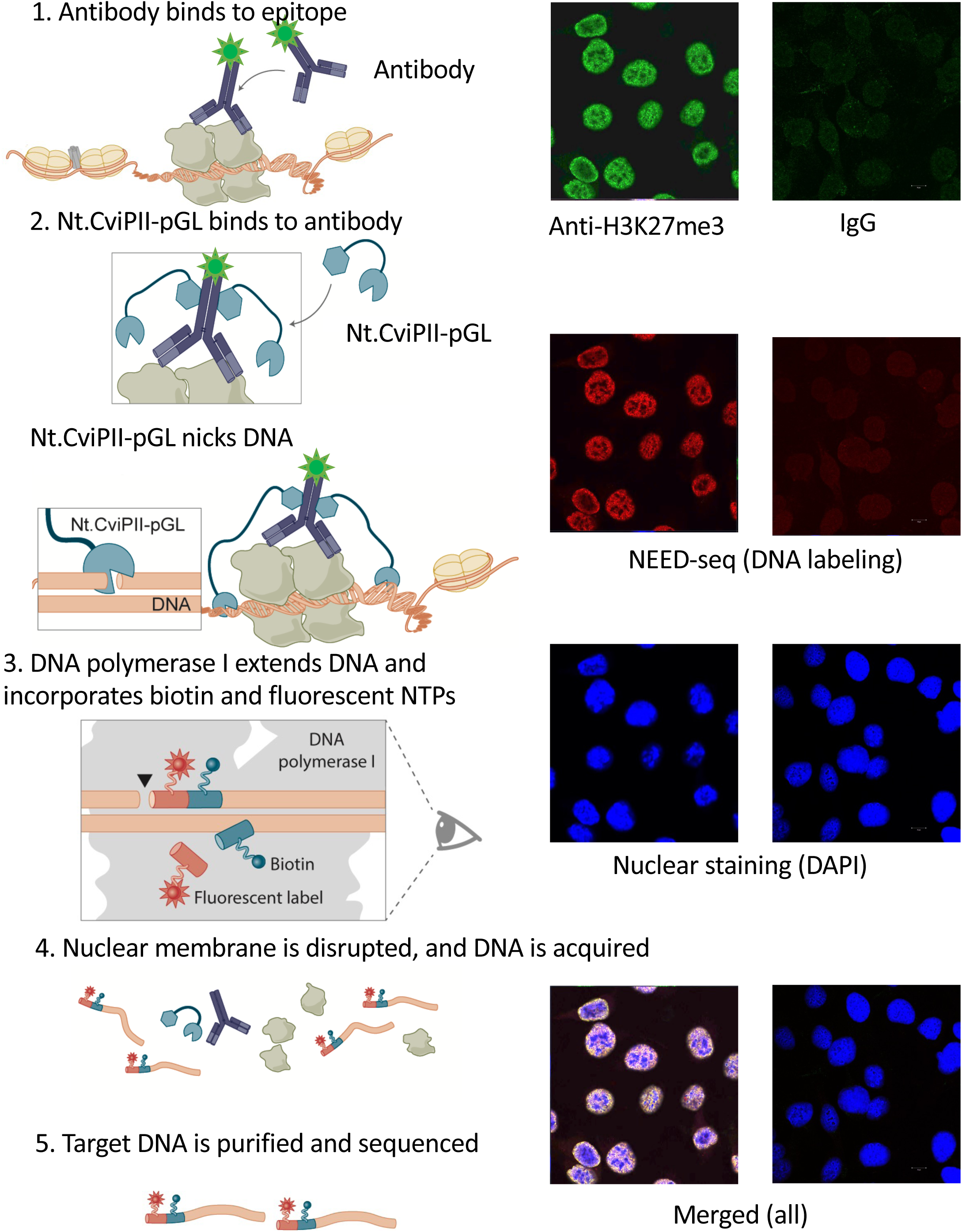
NEED-seq: Schematic diagram depicting the steps of NEED-seq (left panel). Example of NEED-seq images of HT1080 cells using confocal microscopy. Green color represents binding of Alexa488 conjugated H3K27me3 antibody to the epitope, and red shows DNA labeling during polymerase extension using Texas-Red conjugated dATP in the dNTPs mixture (right panel). IgG is the control antibody for NEED-seq.

### NEED-seq

In principle, NEED-seq is a 3-step process. (1) Epitope specific antibody is used for targeting and marking the protein bound to DNA with a fluorophore. To accomplish this, fluorophore conjugated primary or secondary antibody can be used. (2) Nt.CviPII-pGL fusion binds to the antibody and non-specific fusions are washed away. (3) Nicks and nick-translation on the DNA is initiated upon addition of Mg^2+^ and DNA polymerase I in the presence of biotin and/or fluorophore conjugated dNTPs for both visualizations, and streptavidin bead capture of target DNA for library preparation (Fig. 1, left panels). Indeed, Alexa 488-conjugated H3K27me3 antibody was able to label the heterochromatic regions as green, and Texas-red-dATP incorporated epitope bound chromatin as red, resulting in a yellow merged pattern (Fig. 1, right panels) demonstrating the overlap of epitope and DNA bound regions. Non-specific antibody, IgG alone showed no signal in antibody staining or NEED-seq mediated DNA labeling conforming specificity of NEED-seq requires antigen specific antibody (Fig. 1, left panels).

### NEED-seq of histone PTMs

Nicking enzyme fusions, Nt.CviPII-pGL, use DNA as substrate similar to Nt.CviPII alone (Fig. S2). These enzymes also can tether to epitope bound antibody via protein G and L fused domain (Fig. 1). To make the NEED-seq reaction specific to antibody bound epitope, non-specific DNA bound fusion enzymes must be removed by buffer washing steps. We validated NEED-seq for other histone PTMs encompassing both activation marks (H3K4me1, H3K27ac, H3K4me3 and H4K20me1), and repressive marks (H3K9me3, H2AK119ub, H4K20me3 and H3K27me3) using HT1080 cells. Indeed, most of the tested antibodies displayed good signal to noise ratio across chromatin regions. The histone PTM were inversely presented in the genome i.e. activation mark rich chromatin were poor with inactivation marks and vice versa (Fig. 2A). Also, high Pearson’s correlation (r=0.98-0.99) between two replicates (rep1 and 2) was observed for the all the histone PTMs (Fig. 2B). Additionally, NEED-seq exhibited higher fraction reads in peaks (FRiP) score, ranging between 0.2 to 0.37 (Table S1). These results confirm the versatility and reproducibility of NEED-seq in interrogating histone PTMs across different cell lines.

**Figure 2:**
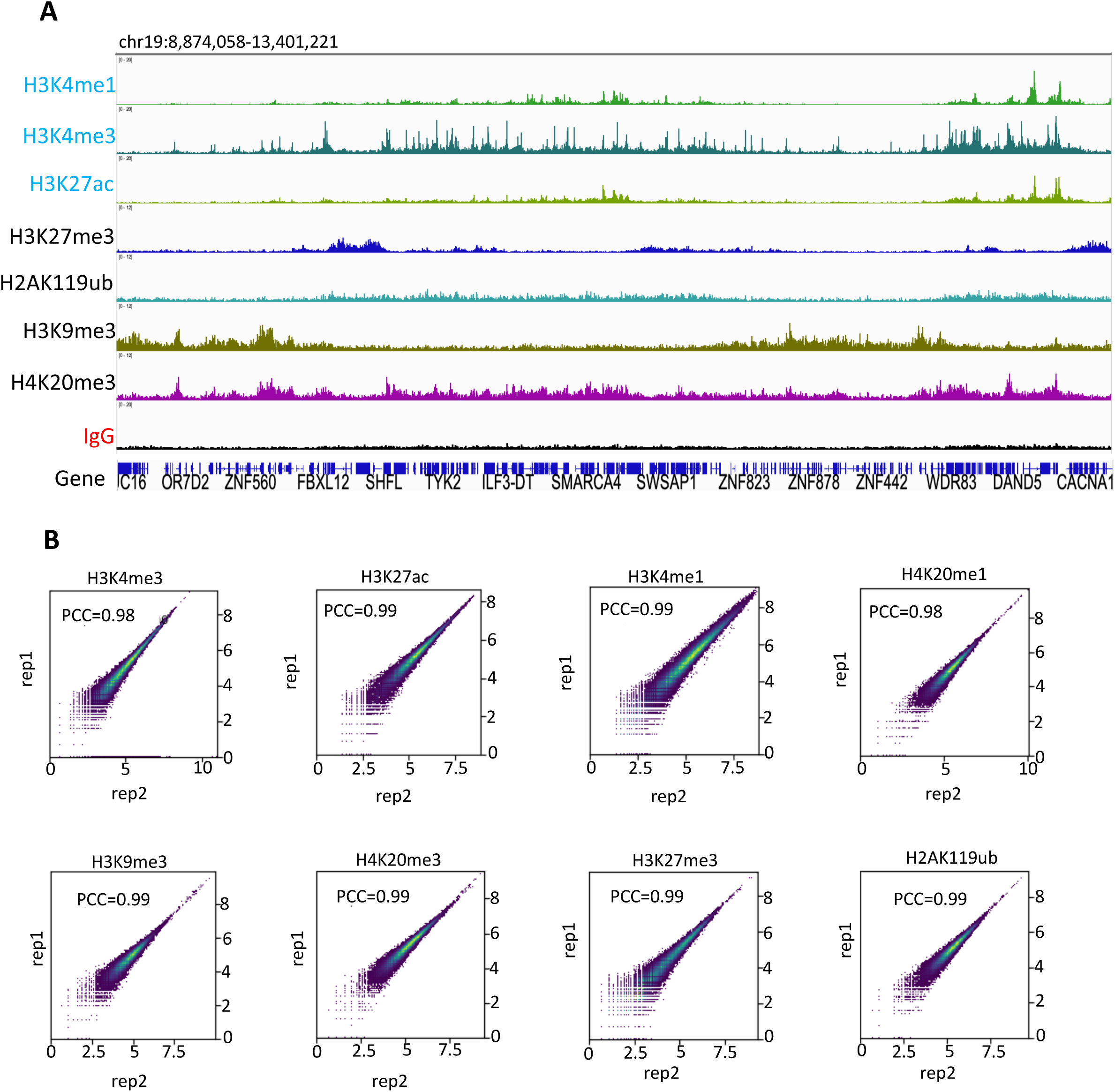
NEED-seq of histone post translational modification in cells. (**A)**. Representative IGV genomic track for different histone marks along with control IgG in HT1080 cell line. **(B)** Pearson Correlation based scatterplots for different histone marks between sequencing replicates of HT1080 cell line. Histone PTM marks are indicated on the top.

### NEED-seq of transcription factors

Since NEED-seq was able to profile histone PTMs in genome-wide, we applied it to low abundance epitope such as transcription factors (TF) bound DNA profiling. We profiled, c-Jun, c-Fos, RB1, p53 genome-wide using specific antibodies along with control IgG. Since c-Jun dimerizes with itself and with c-Fos, forming (c-Jun homodimers and c-Jun-c-Fos heterodimer) complexes of different DNA binding affinities, we hypothesized that we would observe similar NEED-seq patterns for either of them. Indeed, genome-wide chromatin bound pattern for c-Jun and c-Fos were very similar. Both RB1 and p53 displayed unique genome-wide binding pattern with some common peaks with c-Jun and c-Fos (Fig. 3A). This led us to believe that these transcription factors may be in a complex during cellular development. We performed STRING (Search Tool for the Retrieval of Interacting Genes/Proteins) for predicting protein–protein interactions. In fact, c-Jun, c-Fos, RB1 and p53 interact in certain locations in the chromatin (Fig. 3B).

**Figure 3.**
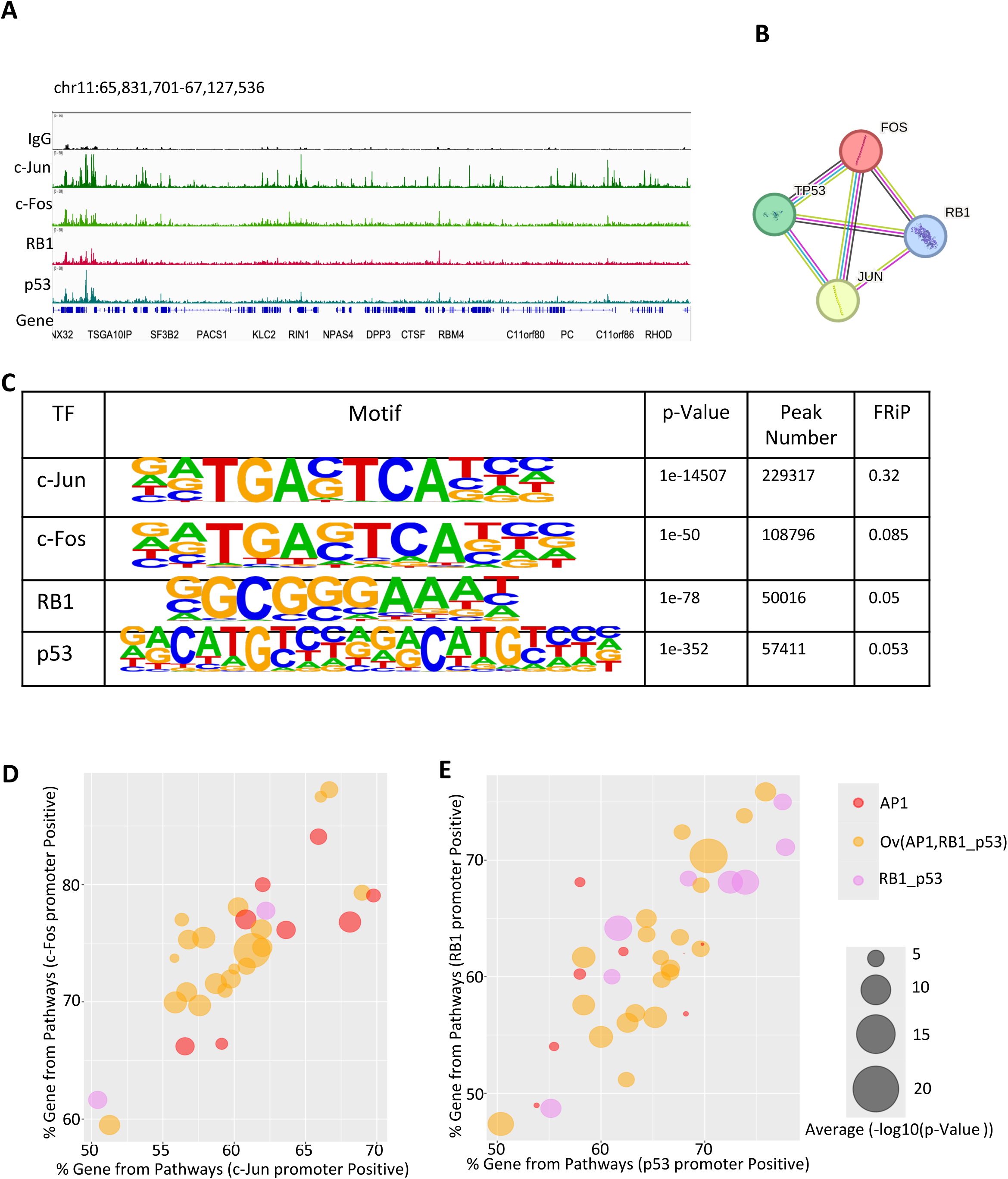
NEED-seq of transcription factor in cells. (**A)**. Representative IGV genomic track for different transcription factors (c-Jun, c-Fos, RB1 and p53) along with control IgG in HT1080 cell line. **(B)** STRING (v 2.0) diagram demonstrating connectivity between c-Jun, c-Fos, RB1 and p53 transcription factors. **(C)** HOMER analysis for transcription factor binding site consensus along with p-values, genome-wide peak numbers and FRiP score. (**D**) Bubble plots comparing gene percentage from pathways between c-Jun promoter positive and c-Fos promoter positive NEED-seq results. AP1 specific (red), overlap of AP1, RB1 and 53 (orange), and RB1 and p53 common (violet) are shown. (**E**) Bubble plots, similar to D, comparing p53 promoter positive and RB1 promoter positive genes percentage derived from associated pathways are shown. Please note that the size of the bubble indicates significance of association.

To unequivocally establish c-Jun, c-Fos, RB1 and p53 are indeed on their respective binding sites, we performed HOMER motif analysis using the BED regions. The known motifs of c-Jun, c-Fos matched with AP1 conserved motif with high p-values of 1e-14507 and 1e-50 respectively with ∼220k and ∼110k specifically identified peaks. These results clearly demonstrate c-Jun, in combination with protein c-Fos, forms the AP-1 early response transcription factor, as shown previously (acc number # MA0099.3). Similarly, p53 known motifs showed similarity with publicly available p53 motif (p-value = 1e-352) with ∼60k identified peaks. For RB1 bound motifs, there was no publicly available motif. However, RB1 is known to bind E2F family genes (Engeland, 2022). We used the ∼50k RB1 bound peaks and identified E2F1 motif with p-value 1e-78. These high p-values demonstrate efficacy of the TF profiling using NEED-seq (Fig. 3C).

Next, c-Jun, c-Fos, RB1 and p53 bound promoter of protein coding genes were identified. Set of genes with bound transcription factor were analyzed for associated pathways using Wikipathways database with stringent criteria (>/=10% gene association and p-value<=0.05). 72 common pathways from c-Jun and c-Fos specific list were then ranked based on average (-log10 (p-value)). Similarly, 149 common pathways from RB1 and p53 specific lists were also ranked. 21 pathways were demonstrated as common pathways between top 30 pathways from the above lists (Table S2; Fig. 3D, E). Bubble plots showed distributions of promoter positive genes where size of the bubbles demonstrated rate of significance. Indeed, c-Jun and c-Fos positive genes exhibiting higher rate of significance for pathways associated exclusively with AP1 and overlapping with RB1 and p53 (RB1_p53) whereas pathways associated only with RB1_p53 exhibited low rate of significance (Fig. 3D). Similarly, RB1_p53 positive genes also exhibiting higher rate of significance for pathways associated exclusively with RB1_p53 and overlapping with AP1 where only AP1 associated pathways were showing extremely low rate of significance (Fig. 3E). Interestingly, pathways uniquely associated AP1 were mostly associated with growth mechanisms e.g., Brain Derived Neurotrophic Factor (BDNF) Signaling Pathway whereas RB1_p53 unique pathways were associated with regulation e.g., DNA damage response, miRNA regulation (Table S2). Those results are consistent with AP-1 transcription factors mediating BDNF positive feedback loop (Tuvikene et al., 2016) as well as RB1_p53 pathway activation during DNA damage and its regulation of miRNA expression (Engeland, 2022; Zhou et al., 2023).

### NEED-seq identify common as well as unique set of chromatins bound epitope regions

ChIP-seq, CUT&RUN and CUT&Tag technologies can identify epitope bound chromatin regions. While ChIP-seq and NEED-seq are generally performed on formaldehyde crosslinked cells, CUT&RUN and CUT&Tag are better suited for native cells, although variation of these technologies is used for formaldehyde fixed cells (Henikoff, et al. 2023). To validate the robustness of NEED-seq, we compared the above technologies using two representative histone post-translational modifications (PTMs), one active mark (H3K4me3) and one repressive mark (H3K27me3). Published data for ChIP-seq (ENCODE), CUT&RUN and CUT&Tag of K562 cells were used for comparison with NEED-seq for H3K4me3 and H3K27me3. The Pearson correlation between all the above methods were above 0.72 for H3K4me3, suggesting a high degree of correlation (Fig. S3A). The FRiP score between all the technologies was above 0.2 and was 2X higher for NEED-seq compared to ChIP-seq (0.52 vs. 0.25, Fig. S3B). Peak calling identified 23K, 22K, 15.2K and 12.6K numbers of peaks in NEED-seq, ChIP-seq, CUT&RUN and CUT&Tag for H3K4me3 respectively. Amongst them most (∼ 12.9K) overlapped, and 2.1K-8.3K were unique (Fig. S3C). When the data were visualized using the IGV browser, peaks common to all data sets displayed similar enrichment (Fig. S3D), except CUT&Tag with comparatively lower signal. Since both NEED-seq and ChIP-seq were based on formaldehyde fixation, we also compared their genomic features and peak overlaps using H3K4me3 mark reads and sample downsizing. Indeed, both NEED-seq and ChIP-seq displayed similar genomic features with more promoters being identified in NEED-seq at or below 4 million reads (Fig. S3E, F). Similarly, most of the NEED-seq and ChIP-seq peaks overlapped (Fig. S3G).

In another example, NEED-seq of H3K27me3 was compared to published ChIP-seq (ENCODE), CUT&RUN and CUT&Tag data sets. The Pearson correlation between CUT&RUN or CUT&Tag with NEED-seq remained at 0.67 compared to 0.77 for ChIP-Seq (Fig. S3H). However, the FRiP scores of CUT&RUN, CUT&Tag and ChIP-seq were below 0.1 compared to 0.27 for NEED-seq (Fig. S3I). Further investigation with peak calling identified 251K, 276K, 19K and 17.5K numbers of peaks in NEED-seq, ChIP-seq, CUT&RUN and CUT&Tag for H3K27me3 respectively (Table S3). Low numbers of peaks in both CUT&RUN and CUT&Tag may be attributed to compact heterochromatin environments where the fusion MNase and Tn5 transposon may have difficulty in enzymatic catalysis or transposition. Indeed, the IGV browser showed poor signal to noise in CUT&Tag track (Fig. S3J). Next, we chose to compare the NEED-seq with ChIP-seq since they were based on formaldehyde fixed cells. About 55% of the ChIP-seq peaks were common with NEED-seq where the IGV also looks identical (Fig. S3J, K).

Next, we compared NPAT (nuclear protein, coactivator of histone transcription) transcription factor as an example to compare between CUT&RUN, CUT&Tag data sets derived from fresh cells vs. NEED-seq performed in 4% formaldehyde fixed cells. The Pearson’s correlation between NEED-seq and CUT&RUN, NEED-seq and CUT&Tag remained at 0.78 and 0.87 respectively (Fig. S3L). Similarly, IGV profile showed most of the peaks identified between CUT&RUN, CUT&Tag, NEED-seq are indeed common with major NPAT peaks at the histone genes H3, H4, H2B and H1. Some peaks were uniquely identified by NEED-seq (Fig. S3M). Therefore, with low background and high mapping efficiency, NEED-seq offers insights into epitope bound DNA irrespective of chromatin topology and compactness.

### Comparison between Nt.CviPII-pGL and pAG-MNase in NEED-seq on formaldehyde fixed cells

Although both NEED-seq and CUT&RUN are powerful technique to study protein-DNA interactions in cells using different fusion enzymes, NEED-seq works efficiently in 4% formaldehyde fixed cells unlike CUT&RUN that relies on pAG-MNase and is sensitive to formaldehyde fixation (Miura and Chen, 2020). We therefore compared both techniques side by side using HT1080 nuclei profiled for H3K4me3 and H3K27me3 marks using DNA size selection protocol (see Material and Methods). In one set of experiment where DNA polymerase I step was omitted, Nt.CviPII-pGL and pAG-MNase were incubated at 37°C separately, and the other had pAG-MNase alone at 4°C as recommended by the CUT&RUN protocol. Targeted fragmented DNA induced by either Nt.CviPII-pGL or pAG-MNase were captured using NEBNext Sample Purification Magnetic beads. High molecular weight DNA were removed by using Ampure XP protocol. Spearman correlation between all three conditions for H3K4me3 remained between r=0.89-0.93; and for H3K27me3, r=0.8-0.93 suggesting high degree of similarity (Fig. S4A, B). However, the FRiP score, peak numbers, enrichment plot, and signal to noise ratio was ∼2X higher in NEED-seq samples compared to CUT&RUN (Fig. S4C-H). Taken together, it is apparent that Nt.CviPII-pGL based NEED-seq offers a robust genome-wide chromatin profiling on formaldehyde fixed cells compared to pAG-MNase.

### NEED-seq with low cell numbers and multiple histone PTMs

Since NEED-seq is capable of deciphering DNA bound epitopes in both H3K4me3 and H3K27me3 regions, we tested its versatility by decreasing the cell numbers from 50K to 50 using anti-H3K4me3 antibody. We chose K562 cells, a chronic myelogenous leukemia patient derived cell line since they grow in suspension and are easy to dilute. The Pearson correlation of different cell numbers with 50 cells was the lowest score of 0.82 (Fig. 4A). A similar trend was also observed in FRiP score analysis, 50 cells being the lowest (0.07) with a gradual increase in the higher cell numbers (50K=0.2; Fig. 4B). Furthermore, the NEED-seq library from 50 cells identified 8.2K peaks, with a vast majority of them (7.1K peaks), overlapping with either 500 (14.5K peaks) and 50,000 (17.1K peaks) cells. This trend reflected a 43% decrease of peak number with 50 cells compared to 500 (Fig. 4C). The genomic features also remained consistent despite cell dilutions with most of the H3K4me3 featured on the promoter (Fig. 4D). We further visualized the heat map between 50, 500 and 50,000 cells using peak ± 3kb criteria and observed a decrease in the peak intensities as the cell numbers were reduced (Fig. 4E). This was further confirmed by visualization of selected genomic regions and observed a consistent overlap in enrichment of sequence tags/peaks (Fig. 4F). Next, we performed NEED-seq for H3K4me3 mark using 50K HT1080 cells, a fibrosarcoma cell line, and compared the epitope binding DNA peaks with K562 cells. 45% of the peaks were common between these two cell lines. In IGV, common and divergent peaks were observed confirming cell line specificity (Fig. 4G, H). Indeed, most H3K4me3 peaks matched with HT1080 accessible chromatin peaks as expected.

**Figure 4:**
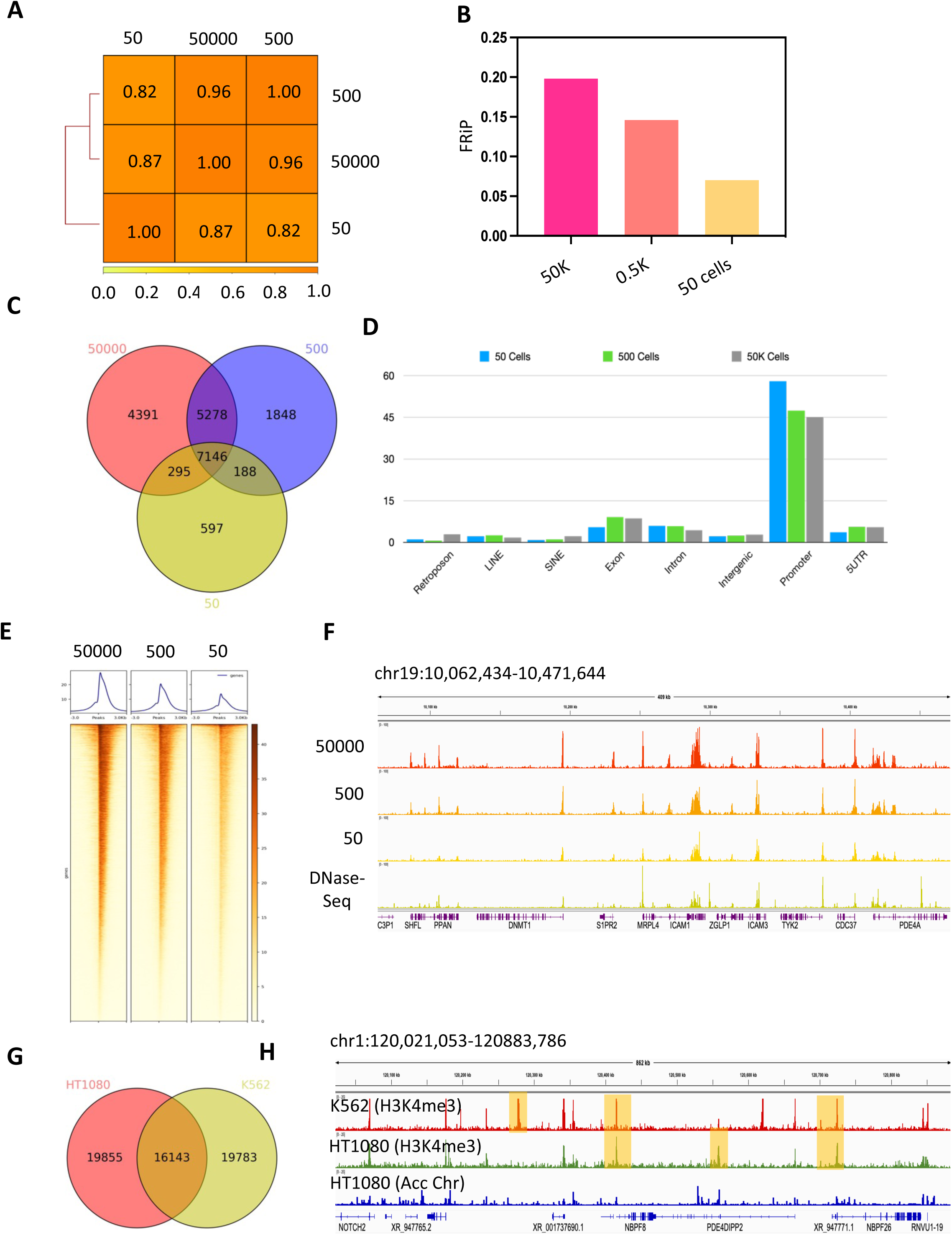
Dynamic range of NEED-seq using H3K4me3 antibody in K562 cells. **(A)** Spearman correlation map on H3K4me3 regions for 50000, 500 and 50 cell numbers. **(B)** FRiP comparison on H3K4me3 regions from 50000, 500 and 50 cell numbers. **(C)** Venn diagram based on overlapping peak regions of H3K4me3 at different cell numbers as indicated. **(D)** Identification of genomic feature specification genome wide from H3K4me3 regions in different cell numbers. **(E)** Genome-wide heatmap showing H3K4me3 at different cell numbers. **(F)** IGV representation of H3K4me3 NEED-seq (50000, 500 and 50 cells) and DNase-seq. **(G)** Venn diagram based on overlapping peaks of H3K4me3 NEED-seq between HT1080 and K562 cell lines. **(H)** IGV representation on H3K4me3 (from K562 and HT1080 cell line) and NicE-seq representing accessible chromatin (Acc Chr, from HT1080) (at data range 0-25) demonstrating unique cell line specific and common peaks, indicated by yellow boxes. Publicly available datasets of NicE-seq (GSM4285582/83) and DNase-Seq (GSM2400371) were used (Table S6).

### NEED-seq can profile compact nuclear structural protein lamin B1 and B2

In immunofluorescent experiment the nuclear envelope encapsulates the genome with lamin B1 (green) and B2 (red) as one of the inner membrane proteins and are in proximity with each other (Fig. 5A; Z-stacks in Fig. S5A). Indeed, lamin B1 and B2 showed an average correlation of 0.79 (N=40; Fig. S5B). To precisely determine the location of lamin B1 and B2 in spatial context we performed LOWESS (Locally Weighted Scatterplot Smoothing) analysis using the same set of 40 nuclei. Immunofluorescence displayed deposition of lamin B1 and B2 in proximity, with lamin B1 towards the nucleus center and lamin B2 projected towards the nuclear membrane (Fig. 5B).

**Figure 5:**
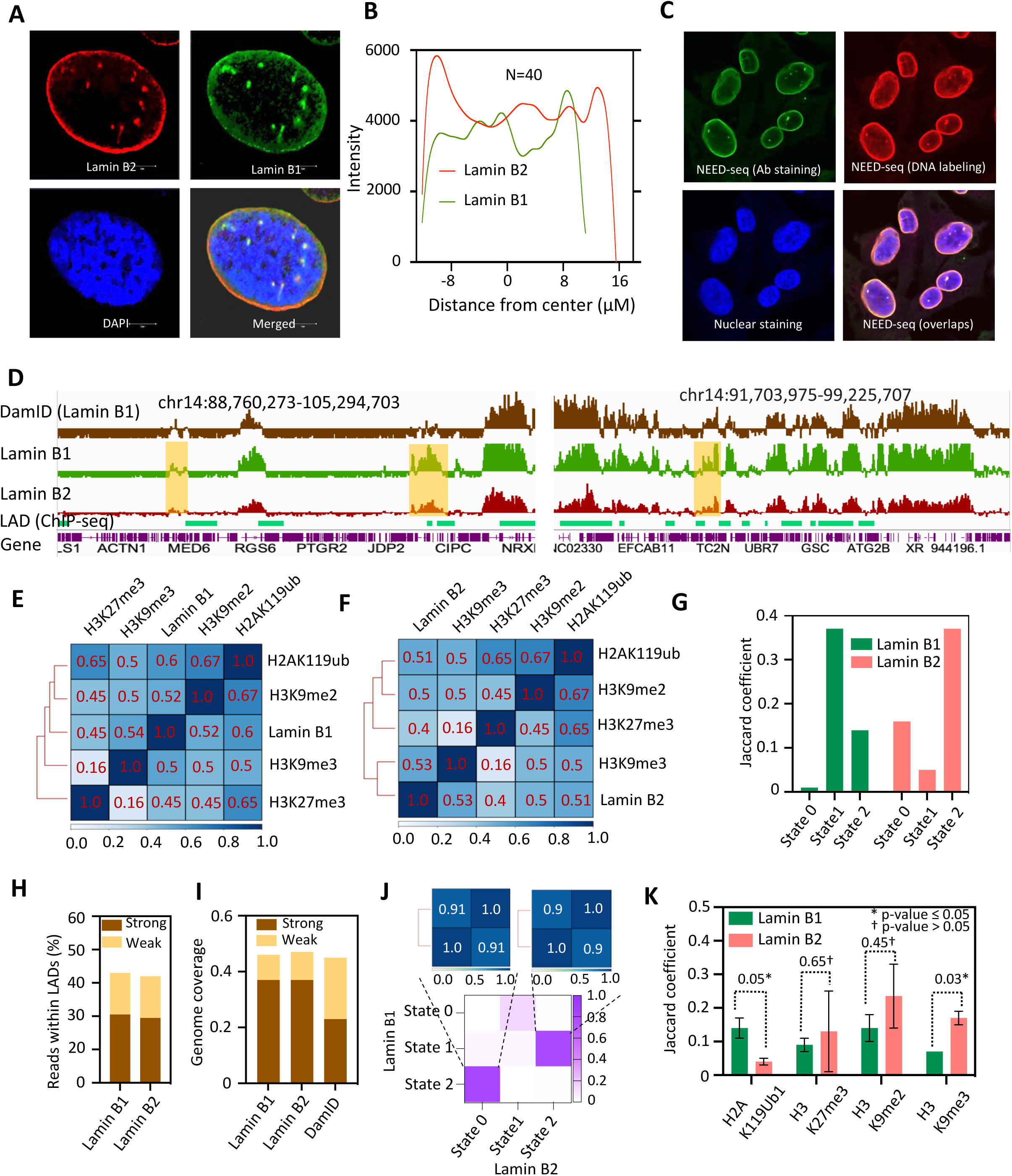
NEED-seq of lamin B1/B2 and association with histone PTMs in HT1080. **(A)** 3D microscopy image (average of 39 Z-stacks) for lamin B1 (green) and lamin B2 (red), where nucleus was stained with DAPI (blue) and co-localization in merged image indicated in yellow. Individual Z-stacks are shown in Fig. S5A **(B)** Intensity vs. distance from center plot for lamin B1 and lamin B2 applying LOWESS is shown. **(C)** NEED-seq microscopy image for anti-lamin B2 Staining (green), DNA labeling (red), and nuclear Staining (blue) is shown. **(D)** IGV visualization for Lamin B1 (DamID, brown), lamin B1/B2 (NEED-seq, green/red) and LAD domains (ChIP-seq, BED regions in pale green) is shown. Yellow shades show lamin B1 and B2 signal differences after performing log2ratio (lamin B1 or B2/IgG). **(E-F)** Spearman correlation with H3K9me3, H3K9me2, H3K27me3, H2AK119ub for lamin B1 and lamin B2 within respective domains are represented. **(G)** Jaccard coefficient (JC) for lamin B1/B2 HMM states (0-2) with LAD ChIP-seq are shown. Defined states for lamin B1 (green) are as follows, State 0, JC <0.1; State 1, JC >0.15; State 2, JC 0.1 to <0.15. Similarly, for lamin B2 (red) State 0, JC 0.1 to <0.15; State 1, JC <0.1; State 2, JC >0.15 were utilized. **(H)** Read enrichment calculation within lamin B1/B2 strong (brown) and weak (yellow) domains are shown. **(I)** Genome coverage comparison between lamin B1/B2 (NEED-seq) and lamin B1 (DamID) strong and weak domains are shown. **(J)** Jaccard similarity calculation between lamin B1 and B2 HMM states (lower panel) and read to read Spearman correlation within strong domains and weak domains (upper panel) are shown. **(K)** Jaccard coefficient score with each histone marks (H3K9me3, H3K9me2, H3K27me3, H2AK119ub) for lamin B1/ B2 strong and weak domains are shown. p-value significance calculation (* p-value ≤ 0.05 and † p-value > 0.05) is indicated.

Since lamin B1, B2 in conjunction with lamin A and C, stabilizes of heterochromatin and chromosomal positioning along the nuclear periphery, these lamina-associated proteins represent compact nuclear structural architecture and are difficult to profile in cross-linked cells. Both DamID and ChIP-seq has been used for lamin B1 profiling. However, detailed lamin B2 bound genomic features are limited. Therefore, we used NEED-seq to profile lamin B2 genome-wide. First, we performed antibody labeling on fixed HT1080 cells displayed lamin B2 presence in nuclear membrane (green), as expected. NEED-seq DNA extension reaction for lamin B2 (with Texas Red-dATP) essentially labeled the same region with red as that of the anti-lamin B2 antibody with ∼0.91 Pearson correlation demonstrating the colocalization of antibody and its bound DNA (Fig. 5C, 2D image). The 3D image conclusively established the NEED-seq labeling in proximity of lamin B2 and its bound DNA (Fig. 5C, 3D images). Next, we proceeded to perform NEED-seq for both lamin B1 and B2 and compared lamin B1 with previously published datasets for validation. The sequence reads of lamin B1 NEED-seq track were compared to previously published DamID. The Pearson correlation between lamin B1 NEED-seq vs. DamID (r=0.72) suggesting strong sequence read correlation (Fig. 5D). We also analyzed the BED files of lamin B1 ChIP-seq (Meuleman et al., 2013), with DamID sand NEED-seq. The UpSet plot of BED files exhibited >50% common regions (Fig. S5E). With these series of comparative analysis, we are confident of NEED-seq profiling of compact chromatin regions.

However, Spearman correlation analysis between lamin B1 and B2 resulted r=0.74, suggesting difference between their DNA binding specificity (Fig. S5F). NEED-seq reads of lamin B1 and B2 were mapped to the human genome (hg38), normalized to SES (Das et al. 2023) (using IgG signal), and represented as log2 (signal/input). Genome browser visual confirmed the presence of discrete lamin B1 and B2 enriched domains consistent with the presence of LADs (Fig. 5D). To determine the differences between lamin B1 and B2, we investigated their association with histone PTMs that would correlate with structure and function. The histone modifications H3K9me2 and H3K9me3 are shown to be enriched in LADs and are often associated with transcriptional repression and heterochromatin formation. Similarly, H3K27me3 and H2AK119ub, also known as Polycomb repressive mark, are often enriched in facultative heterochromatin. We predicted that lamin B1 and B2 NEED-seq identified DNA localizations would therefore correspond to the above heterochromatin marks. We performed NEED-seq using anti-H3K9me2, -H3K9me3, -H3K27me3, -H2AK119ub and compared with lamin B1 and B2 (Fig. 5E, F). As expected, both lamins exhibited high degree of correlation with, H3K9me2 and H3K9me3. However, a small yet significant difference between lamin B1 and B2 with H2AK119ub emerged (lamin B1, r=0.6; lamin B2, r=0.5). Therefore, we segregated lamin B1 and B2 domains (LADs) to three different states using HMMbigwig (github.com/gspracklin/hmm_bigwigs). The states were defined based on their statistical similarity and diversity when compared to LAD-ChIP-seq regions using Jaccard coefficient (JC): no LAD (JC <0.1), weak LAD, JC >0.1-0.15; and strong LAD JC >0.15). Lamin B1 predominately exhibited state 1 (strong LAD) in contrast with state 2 (weak LAD). Similarly, lamin B2 domains were predominately state 2 (strong LAD) and state 0 (weak LAD) (Fig. 5G). In addition, ∼40% of NEED-seq reads were within lamin B1 and B2 strong and weak LADs (Fig. 5H). Since DamID is extensively used for LAD analysis, we compared genome coverage of strong and weak LADs in lamin B1, lamin B2 (NEED-seq), to DamID. For this analysis we used IgG NEED-seq reads for background subtraction from anti-lamin B1/B2 NEED-seq to identify LAD domains utilizing the SES (signaling extraction scaling) method as demonstrated previously (Das et al. 2023). We observed higher percentage of strong signal in NEED-seq compared to DamID demonstrating the robustness of NEED-seq (Fig. 5I). Additionally, we compared different state exhibited by lamin B1 and B2 to correlate association of lamin B1 and B2 states using Pearson’s correlation and observed state 0 of lamin B2 and state 2 of lamin B1 (weak LAD); state 2 of lamin B2 and state 1 of lamin B1 (strong LAD) reads with r=0.9 correlation (Fig. 5J). This suggests lamin bound domains are segregated based on the state and are differentially represented between both lamin B subtypes. Next, we compared lamin B1 and B2 (strong and weak domains) with NEED-seq bed files for H3K9me2, H3K9me3, H3K27me3, and H2AK119ub marks to determine their chromatin association. To our surprise lamin B1 significantly associated with H2AK119ub (p=0.05) and lamin B2 significantly associated with H3K9me3 (p=0.03; Fig. 5K). This was also evident in chromosome wide distribution of H2AK119ub and H3K9me3 between lamin B1 and B2 respectively since they were represented in positive quadrant of individual chromosome (Fig. S6A-D).

### Analysis of lamin B1 and B2 in chromosomal compartment

Since lamin B1 is significantly associated with H2AK119ub and it can control H3K27me3 deposition on chromatin by regulating PRC2 recruitment (Tamburri et al., 2020), we performed ChromHMM analysis for lamin B1 and compared with lamin B2 to decipher functional annotation. As expected, lamin B2 remain associated with constitutive heterochromatin state. However, lamin B1 displayed strong chromHMM state emission with both weak and strong repressed Polycomb (Fig. S7A). Therefore, we hypothesized that lamin B1 and B2 would associate distinctly with chromosomal compartments. To study this association, we performed chromosomal conformation capture experiment (Hi-C). Analysis of the Hi-C data of HT1080 cells showed alignment consistency of 72% between two read tags (Fig. S8A). From the aligned reads, 86% valid interaction were distributed as 77% cis- and 9% trans interactions (Fig. S8B). Next, we transformed interaction map in “obs_exp” matrix (Fig. S8D). Further compartment analysis on interaction matrix accounted for ∼26K A-compartments and ∼27K B-compartments from cis TADs (topologically associated domains) at 50 kbp resolution (Fig. S8C). We first analyzed lamin B specific LAD and their association with chromosomal conformation in HT1080 based on Jaccard coefficient (weak and strong LADs). We observed ∼2.5X association of lamin B1/B2 with B-compared to A-compartment (Fig. 6A). Next, each lamin B1/B2 domains within A/B compartments were analyzed for percentage overlap with DNA methylated regions using HT1080 EM-seq data, since CpG methylation is prevalent in heterochromatic region (Fig. 6B). Approximately ∼35% lamin B1/B2 domains within A-compartments and ∼55% lamin B1/B2 domains within B-compartments were methylated. Although, methyl CpG distribution in lamin B1 domains within B-compartment were lower compared to lamin B2 (Fig. S7B).

**Figure 6:**
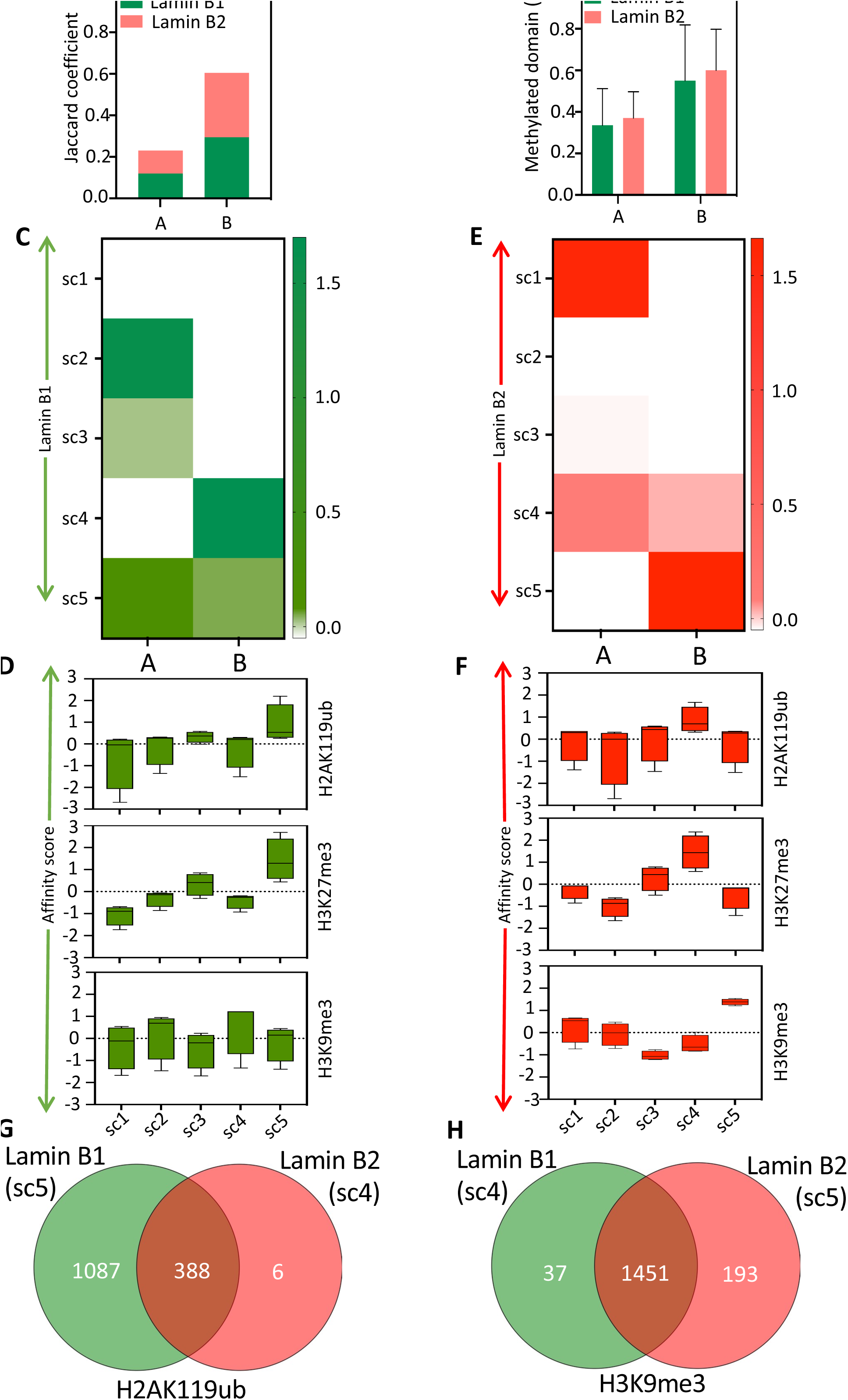
Lamin B1 and B2 in chromosomal compartment and histone portioning analysis. **(A)** Association of lamin B1/B2 with A or B compartment derived from Hi-C based on Jaccard coefficient is shown. **(B)** Percentage of methylated LAD domains within A/B compartments for lamin B1 and B2 is shown. **(C)** Affinity of lamin B1 subcompartments with A/B compartments. **(E)** Similar to C, affinity of lamin B2 subcompartments with A/B compartments. **(D)** Affinity of lamin B1 subcompartments with H2AK119ub (upper), H3K27me3 (middle), and H3K9me3 (lower) and exhibiting the histone partitioning is shown. **(F)** Similar to D, showing affinity of lamin B2 subcompartments with H2AK119ub (upper), H3K27me3 (middle), and H3K9me3 (lower) and exhibiting the histone partitioning is shown. (**G**) Venn diagram of genes H2AK119ub partition with lamin B1/B2 and associated A compartment bound promoters. (**H**) H3K9me3 partition with lamin B1/B2 and associated B compartment bound promoters derived from featured annotation is shown.

Next, we studied the association between LAD identified by NEED-seq with genome-wide chromosome conformation in Hi-C. Therefore, we performed k-means clustering (k = 5) on lamin B1/B2 domains and 1^st^ principal component of Hi-C (PC1) for structural partition to define subcompartments within 25kb bin size. These domains were clustered in five exclusive subcompartments (sc) with lamin B1 and B2 (Fig. 6C, E). Lamin B1 subcompartments sc2 and 4 exhibited strong affinity with A and B compartments (Fig. 6C). Similarly, sc1 and sc5 revealed strong affinity with A and B compartment respectively, for lamin B2 (Fig. 6E). The affinity of subcompartment from lamin B1/B2 is mostly similar with B-compartment. However, lamin B1 exhibits a relatively higher affinity with A compartment compared to lamin B2. Furthermore, lamin B1 sc5 and lamin B2 sc4 also exhibited affinity with both A/B compartments (Fig. 6C, E).

### Lamin B1 and B2 subcompartments show distinct heterochromatin partitioning and biological processes

Since heterochromatin marks are associated with lamin B1/B2, we hypothesized that lamin B subcompartments may have differential affinity for specific heterochromatic mark. We analyzed H3K9me3, H3K27me3, and H2AK119ub with each subcompartment of lamin B1/B2 (Fig. 6D, F). Indeed, Polycomb marks (H2AK119ub, H3K27me3) showed stronger affinity score with lamin B1 compared to lamin B2 (Fig. 6D, F). Similarly, constitutive heterochromatin mark (H3K9me3) showed stronger affinity with lamin B2 compared to lamin B1 (Fig. 6D, F). Individual subcompartments analysis revealed, sc3 and sc5 of lamin B1 and sc4 of lamin B2 exhibited high affinity with Polycomb marks (Fig. 6D, F). We also identified that only lamin B1 sc4 exhibited low affinity score for heterochromatin mark contrary to lamin B2 sc5 that exhibited high affinity for H3K9me3 (Fig. 6D, F).

Since Polycomb mark H2AK119ub exhibited stronger affinity in lamin B1 sc3 and sc5 as well as in lamin B2 sc4, we identified the promoters and unique genes associated with those subcompartments of lamin B. We observed ∼1000 unique genes to lamin B1 in contrast with a handful (∼6) for lamin B2. This was indeed expected since lamin B2 binds to gene poor regions (Fig. 6G). We next performed gene ontology analysis of the lamin B1 unique, as well as lamin B1 and B2 common genes to determine their association with the biological processes. The top four biological processes represented neuronal generation, morphogenesis, development, and synaptic transmission specific to lamin B1 binding. Similar analyses were performed with strong affinity partitions for H3K9me3 in lamin B1 sc4 and lamin B2 sc5. As expected, majority of the genes from these subcompartments (∼1450) were common, 193 unique to lamin B2, and 37 unique to lamin B1 (Fig. 6H). Among the common genes representing biological processes sensory perception on smell and chemicals remain on top followed by cell junction assembly (Table S4). When the identified genes from Fig 6G and H with H2AK119ub or H3K9me3 were mapped to RNA expression profile, genes in sc4 and 5 predominately segregated to either A or B compartments with either higher or lower gene expression (Fig. S7C).

### NEED-seq of lamin B1 and B2 show conserved and divergent domains with defined polymer structure

We next focused on Polycomb (H2AK119ub) and constitutive heterochromatin (H3K9me3) mark distribution in subcompartmental partitioning in chromatin bound regions. For this analysis we merged subcompartment domains and color coded them based on H3K9me3 and H2AK119ub bound vs. unbound. As expected, lamin B2 exhibited higher enrichment within H3K9me3 compared to lamin B1 and vice versa (Fig. 7A). Further metagene analysis of H2AK119ub (lamin B1, sc5; lamin B2, sc4) and H3K9me3 (lamin B1, sc4; lamin B2, sc5) confirmed the IGV profiles (Fig. 7B, C). Next lamin B1/B2 subcompartment histone partitions were analyzed for CpG methylation using HT1080 EM-seq. Partitions for Polycomb marks exhibited hypomethylation (∼15%-20% per CpG methylation) whereas partitions for H3K9me3 exhibit hypermethylation (∼25%-55% per CpG methylation) (Fig. 7D). Therefore, lamin B2 bound DNA is hypermethylated compared to lamin B1.

**Figure 7:**
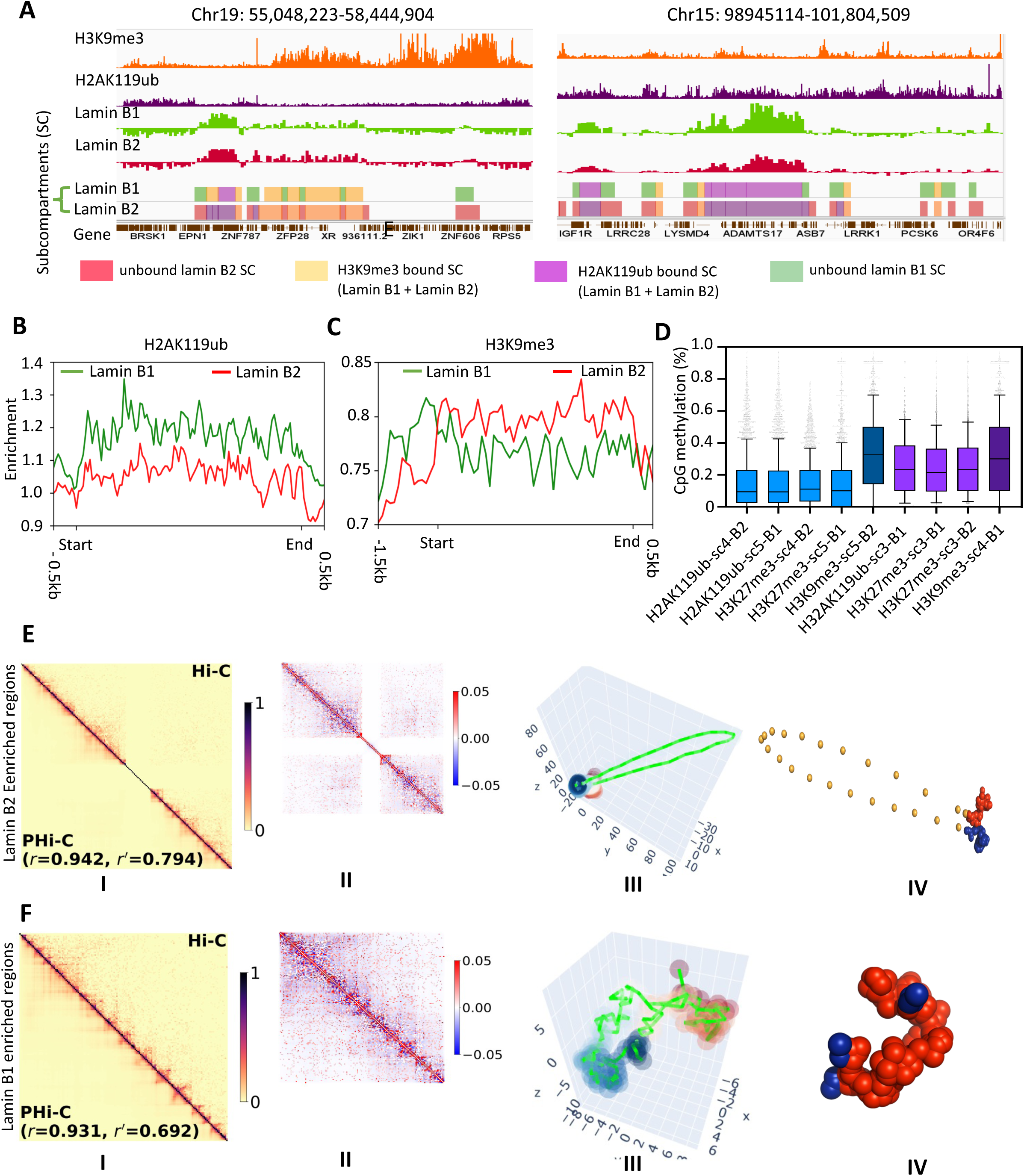
NEED-seq of lamin B1 and B2 and displaying conserved and divergent domains with polymer modeling. **(A)** H3K9me3 (left) and H2AK119ub (right) enriched regions with lamin B1 and B2 based partitioning. Unbound lamin B2 subcompartments (red), H3K9me3 bound subcompartments (yellow), H2AK119ub bound subcompartments (violet), and unbound Lamin B1 subcompartments (green) are shown. **(B-C)** Metagene plots for H2AK119ub (left) H3K9me3 (right) at lamin B1/B2 subcompartments are shown. **(D)** CpG methylation percentage of lamin B1 and B2 subcompartments within polycomb repressive complex and constitutive heterochromatin with high affinity (shades of blue) and low affinity partitions (shades of violet) are shown. **(E-F)** Polymer predication profiles for H3K9me3 (top) and H2AK119ub (bottom) enriched regions using PyMOL is shown. I. Pearson correlation between target regions and genome wide Hi-C. r and r’ represents PCC before and after noise reduction. II. Normalized Hi-C interaction maps. III. Distribution of polymer network model throughout the interaction space. IV. Predicted polymer structure from PyMOL.

Next, we predicted polymer structures from H3K9me3 bound lamin B2 (Chr19: 49,573,757-58,617,617) and H2AK119ub bound lamin B1 (Chr15: 94,123,299-101,991,190) enriched regions from Hi-C using a 4D simulation method, PHi-C2 (Polymer dynamics deciphered from Hi-C data; Shinkai et al., 2020). Overall, both the regions maintain high correlations with genome-wide chromosomes conformations (r=0.94, r’=0.79 and r=0.93, r’=0.69 respectively) (Fig. 7E-I, F-I) However, the chromosomal interaction profiles within lamin B2 enriched regions were segregated in two specific locally conserved domains (Fig. 7E-II). Whereas the chromosomal interaction profiles within lamin B1 enriched regions were scattered genome-wide (Fig. 7F-II). Predicted polymer structures exhibited interaction pattern within lamin B2 enriched regions to be more locally conserved compared to lamin B1 enriched regions (Fig. 7E-III, F-III). A PyMOL representation of both lamin B2 and B1 displayed the predicted polymeric structure of respective locus (Fig. 7E-IV, F-IV). Taken together, predicted polymer structures agree with compactness of the constitutive heterochromatin associated H3K9me3-lamin B2, and the flexible structural conformation of H2AK119ub-lamin B1 can represent facultative heterochromatin in B and A compartments respectively (Graphical Abstract).

## Discussion

Functional genomics studies are used for identifying genome-wide DNA binding sites for transcription factors, other proteins, and post-translational modifications of histones. This has provided crucial clues to regions of transcriptional regulation and chromatin architecture of cells. ChIP-seq, CUT&RUN and their derivative techniques have help propel these studies. Standard ChIP-seq relies on 1% formaldehyde fixation often leading to DNA damage and poly ADP-ribosylation (Beneke et al., 2012; Vishnu et al., 2021). Similarly, CUT&RUN works efficiently on unfixed cells and perform poorly on weakly bound epitope. In contrast with ChIP-seq and CUT&RUN, NEED-seq is a simple method for chromatin features profiling in cells with available antibodies using 4% formaldehyde crosslinking thus fixing the weakly bound transcription factors or proteins profiling. Furthermore, it represses both DNA damage and poly ADP-ribosylation in cells that may otherwise introduce artifacts or hinder antibody targeting (Vishnu et al., 2021). Using the NEED-seq, we have demonstrated its applicability to genome wide histone PTMs (active and inactive marks) and lamin B1 and B2 occupancy. Furthermore, epitope bound DNA can be dual labeled with fluorophore and molecular handle, such as biotin, in a controlled manner without the risk of non-specific labeling. The labelled DNAs are captured on magnetic beads for library preparation or other downstream application to reveal visual and functional genomics. Indeed, the comparison between NEED-seq, ChIP-seq, CUT&RUN and CUT&Tag revealed a large percentage of identified sites were common amongst them. This suggests that NEED-seq is comparable in identifying the genomic features with additional versatility of visual detection, that other technologies currently do not offer.

When we performed NEED-seq of H3K4me3 with gradual decrease in the cell numbers to 50-50,000, most of the identified peaks remained in the dataset with a large fraction being common. A similar number of 100 cells for CUT&RUN against prevalent histone marks has been reported (Skene et al., 2018). NEED-seq was superior to ChIP-seq due to lower cell numbers and fewer steps, although single cell ChIP-seq can be performed with unique barcodes. In this protocol, the single cells are first isolated into droplets containing lysis buffer and MNase and then fused to a droplet carrying specific barcoded oligonucleotides with sequencing adaptors, and restriction sites. DNA ligase was also included in the droplet to complete the tagging process. Furthermore, carrier chromatins must be introduced into the pooled droplets before chromatin immunoprecipitation. Library preparation is carried out according to ChIP-seq procedures before sequencing. Indeed, a high proportion of reads aligns to genomic positions enriched in both bulk ChIP-seq assays and aggregated chromatin profiles from 200 single-cell for both ES and MEF cell lines, providing evidence that single-cell data are informative (Rotem et al., 2015).

NEED-seq of compact structural lamin B1 and B2 demonstrated their visual presence in the nuclear membrane and lamin associated domains (LADs) in fixed cells. This is in contrast with DamID and ChIP-seq that do not provide information on visualization of LADs in the nucleus. Although in DamID the visualization can be achieved by fluorescence in situ hybridization (FISH) using LAD sequences specific FISH probes or a m6A binding protein fused with GFP (Kind et al., 2013). Despite methodological differences, lamin B1 NEED-seq LADs significantly overlaps with DamID and ChIP-seq (Fig. S5E). Another method, pA-DamID is an alternative implementation of protein A-based enzyme targeting to study protein–DNA interactions has been applied to study lamin. This method requires *in situ* DNA methylation reaction as well as use of N6A-tracer for visualization (vanSchaik et al., 2020).

Both lamin B1 and B2 bound chromatin are enriched with histone H3K9me2 and me3 of constitutive heterochromatin and possess low expressed and repressed genes (Lund et al., 2014; Guelen et al., 2008). Here we show that lamin B1 was comparatively more enriched for Polycomb mark (H2AK119ub and H3K27me3) compared to lamin B2. These marks are propagated by PRC1 and PRC2 and an integral part of gene repression. Indeed, loss of H2AK119ub induced a rapid displacement of PRC2 activity leading to loss of H3K27me3 deposition affecting gene expression. Hypothetically, the excess Polycomb repressive deubiquitinase (PR-DUB) complex would enzymatically remove the ubiquitin moiety from H2AK119ub on the nucleosome and affect nuclear architecture and/or gene expression. It has been reported that a fraction of lamin B can interact with expressed genes during epithelial to mesenchymal transition (Pascual-reguant et al., 2018). Interestingly, reader, writer, and eraser of H2AK119ub are recurrently mutated in myeloid disorders such as AML, including ASXL1, DNMT3A, BCOR and BCORL1 (Döhner et al., 2017).

Instead of being evenly distributed along the entire nuclear periphery, lamin B1 and B2 were located asymmetrically in one segment of the nucleus. Often lamin B2 was located entirely within the large, solitary nuclear bleb (Fig. 5A) which was previously observed in neurons (Coffinier et al., 2011). Measuring their signal from nucleus center one could see both lamin B1 and B2 can position themselves either at the inner or the periphery of the nuclear membrane. Lamin B1 and B2 remain associated with the nuclear membranes absorbed into the ER due to their farnesyl anchors. However, other compounding factors such as cell cycle stage and gene expression can’t be excluded for such observation.

By utilizing chromHMM analysis for lamin B1 and lamin B2, we discovered their association with both weak and strong repressed Polycomb. However, lamin B2 remained strongly associated with heterochromatin. Despite, high degree of sequence similarity between lamins B1 and B2 they are not functionally redundant. Knockout studies of the respective genes *Lmnb2-/-* and *Lmnb1-/-* in mouse demonstrated that both B-type lamins are crucial for neuronal development and migration (Kim et al., 2011;). Knockout *Lmnb2-/-* or *Lmnb1-/-* mice died shortly after birth with neuronal migration abnormalities in the cerebral cortex, and a small cerebellum devoid of sulci. This observation was further substantiated by our gene ontology analysis of A compartment bound lamin B1 and B2 with strong affinity for H2AK119ub. Indeed, lamin B1 bound genes were mostly associated with neuronal developmental pathways and lamin B2 in sensory perception and cell-cell junction (Table S4). This would suggest that both lamin Bs functional pathway events contribute to cell division and structural integrity of the nuclear lamina. In another sudy it was demonstrated that lamin B1 is essential for mature neuron specific gene expression (Gigante et al., 2017). Thus, we hypothesize that when lamin Bs are impaired/abolished, the aberrant cell division control may affect cerebellum development, and aberrant neuronal pathways may allow the forces generated by the cytoplasmic motors simply stretching out the nucleus rather than translocating it into territory at the leading edge of the cell. Taken together, lamin B1 and B2 bound chromatin region are not only structural, but they also participate in cell fate and development.

## Materials and Methods

### NEED-seq for adherent cells

HT1080 and HCT116 cells were grown in 6 well plates until 70-80% confluency. After one wash with 1X PBS (GIBCO # 770011-044), cells were crosslinked in 1X PBS with 4% formaldehyde (Thermo Fisher Scientific # 28908) for 10 min at room temperature (RT). Crosslink was quenched using 0.25 M Glycine for 5 min at RT. Cells were de-crosslinked overnight at 58°C in 1X PBS. Cytoplasm was extracted for 10-30 min on ice using cytosolic buffer (15 mM Tris-HCl, pH 7.5, 5 mM MgCl_2_, 60 mM KCl, 0.5 mM DTT, 15 mM NaCl, 300 mM sucrose, and 1% NP40) with occasional agitation. After 1X PBS wash, cells were incubated with 1X PBS / 5% BSA / 0.3% Triton™ X-100 for 1 hr at RT with gentle rocking. Primary antibodies were added (1/400 dilution per well) and incubated overnight at 4°C with gentle rocking. After 3 washes with 1X T-PBS (PBS + 0.1% Tween 20) for 5 min at RT, secondary antibodies were added, Alexa Fluor® 488 conjugated goat anti-Mouse IgG (H+L) Superclonal™ or goat anti-Rabbit IgG (H+L) Superclonal™ (corresponding to species of primary antibody; Thermo Fisher # A-28175 and A-27034 respectively) at a dilution of 1:1000 in PBS / 0.3% Triton™ X-100/ 5% BSA for 1 hr at RT with gentle rocking. After 3 washes with 1X T-PBS for 5 min at RT, cells were incubated for 1 hr at RT with 1 µl of Nt.CviPII-pGL (NEB, 0.5 units) in 800 µl binding buffer per well (20 mM potassium phosphate pH 7.0, 20 mM NaCl, 0.3% Triton™ X-100). Plates were then washed 3 times with washing buffer (20 mM potassium phosphate buffer pH 7.0, 500 mM NaCl, 0.3% Triton™ X-100) for 15 min at RT. Cells were incubated with 10 units of DNA Polymerase I (NEB # M0209S), 5 μM of dNTP mix including Biotin-14-dCTP (Thermo Fisher # 19518018), 5-methyl-dCTP (NEB # N0356S), dATP, dTTP and dGTP for 1 hr at 37°C in 800 µl of 1X NEBuffer 2 per well to allow nick translation to occur. Plates were then washed 3 times with washing buffer (20 mM potassium phosphate buffer, pH 7.0, 500 mM NaCl, 0.3% Triton™ X-100) for 15 min at RT with an additional wash with 1X PBS. Cells were then covered with 100 µl of 1X PBS and DNA was extracted using Monarch genomic DNA purification kit (NEB # T3010S) according to manufacturer’s recommendations. After DNA purification, 100-500 ng of genomic DNA was fragmented using 2 units of Nt.CviPII (NEB # R0626S) in 500 µl of 1X NEBuffer 2 at 37°C for 4 hrs to overnight. Nt.CviPII was inactivated at 65°C for 20 min. NaCl 5M stock was added to reach 1 M final concentration along with 30 µl of Streptavidin magnetic beads (NEB # S1420S) for 2 hrs with end-over-end mixing at 4°C to capture biotinylated DNA fragments. Streptavidin bound DNA beads were washed twice with 1X TE buffer supplemented with 2 M NaCl and 0.05 % Triton™ X-100. A third wash was performed with 1X TE buffer. Beads were resuspended in 50 µl of TE buffer and libraries were made on beads using NEBNext Ultra™ II DNA Library Prep Kit for Illumina (NEB # E7645S) according to manufacturer’s protocol. 8 to 12 PCR cycles were used to amplify barcoded DNA. 1 nM of size selected DNA library was sequenced on Novaseq 6000. Details of antibodies are provided in Table S5.

To visualize NEED-seq using microscopy, cells were grown on coverslips and 0.5 μM of fluorescently labeled dATP (Perkin Elmer # NEL471001EA) was incorporated in the dNTP mix using NEED-seq protocol as described above. After the last wash with 1X PBS, coverslips were dried and mounted on slides using SlowFade® Gold Antifade Mountant with DAPI (Thermo Fisher # S36938). Alexa Fluor® 488 conjugated secondary antibodies, Goat anti-Mouse IgG (H+L) Superclonal™ or Goat anti-Rabbit IgG (H+L) Superclonal™ and Texas-Red-5-dATP were detected using 458, 488, 514 nm multiline Argon laser and 561 nm DPSS laser, respectively. Images were captured using a confocal microscope (LSM 880, Zeiss).

### NEED-seq for suspension cells

NEED-seq protocol described above was used with some modifications. Briefly, K562 crosslinked cells with 4% formaldehyde were decrosslinked overnight at 58°C in 1X PBS. Cytoplasmic extraction was performed for 30 min with end-over-end mixing at 4°C. After 300 x g centrifugation at 4°C for 5 min, cell pellet was washed once with 1X PBS and nuclei number was calculated using Luna automated cell counter. Nuclei resuspended in 1X PBS were then captured by adding 10 µl of preactivated Concanavalin A magnetic beads (Cell Signaling Technology # 93569) for 10 min with end-over-end mixing at RT. Bead-bound nuclei were incubated with 1X PBS including 0.3 % Triton X-100 and 5% BSA for 1 hr at RT. Primary antibodies were added (1/400 final dilution) and incubated overnight with end-over-end mixing at 4°C. After 3 washes with 1X T-PBS for 5 min at RT, blocking buffer was added with anti-mouse IgG or anti-rabbit IgG coupled with Alexa 488 for 1h at RT with gentle rocking. After 3 washes with 1X T-PBS for 5 min at RT, beads were resuspended in 1 ml binding buffer (as described above) including 1 μl of fusion enzyme Nt.CviPII-pGL (NEB) for 1 hr at RT. Beads were then washed 3 times with washing buffer (as described above) for 15 min at RT. After 1 wash with 1X PBS, beads were incubated with 10 units of DNA Polymerase I, 5 μM of dNTP mix including Biotin-14-dCTP, 5-methyl-dCTP, dATP, dTTP and dGTP for 1 hr at 37°C in 800 µl of 1X NEBuffer 2 with end-over-end mixing. After 3 washes with washing buffer for 15 min at RT, beads were resuspended in 50 mM Tris-Cl, pH 7.5, 1 % SDS, 200 mM NaCl and 2 µl of proteinase K (NEB # P8107S) and incubated overnight at 65°C to lyse nuclei bound to the beads and DNA was extracted using phenol-chloroform and precipitated using glycogen. 200 ng of DNA was fragmented using Nt.CviPII and used to 25 libraries as described above. For low number of cells, barcoded DNA was amplified using ∼18 PCR cycles. To determine the dynamic range of NEED-seq, cells were serially diluted after the labeling reaction. 1 nM of size selected DNA library was sequenced on Novaseq 6000. Details of antibodies are given in Table S5.

### NEED-seq biotin free (DNA size selection)

In this modified NEED-seq method, HT1080 cells were grown on 6 well plate to 80% confluency. Cells were incubated for 10 min at 4°C using CSK buffer (10 mM PIPES pH 6.8, 300 mM sucrose, 100 mM NaCl, 1.5 mM MgCl2 and 0.5 % Triton X-100) followed by crosslinking with 4% formaldehyde for 10 min at RT. Glycine was used to quench formaldehyde for 5 min at RT (final concentration 0.25M). After 1X PBS wash, blocking buffer was added to cells for 1h at RT (5% BSA, 1X PBS and 0.3% Triton X-100). Different antibodies (see Table S5) were then added at dilutions recommended by the manufacturer and incubated with cells for overnight at 4°C. After 3 washes with T-PBS (PBS with 0.1 % Tween 20) for 5 min at RT, blocking buffer was added with anti-mouse IgG or anti-rabbit IgG coupled with Alexa 488 for 1h at RT. After 3 washes with T-PBS for 5 min at RT, NEED-seq binding buffer (20 mM KPO4, 150 mM NaCl and 0.3% TritonX-100) was added along with Nt.CviPII-pGL enzyme (1 unit per well) for 1h at 37°C or pAG-Mnase (1.5 μl per well, Cell Signaling Technology # 40366S). After 3 washes with NEED-seq washing buffer (20 mM KPO4, 500 mM NaCl and 1% TritonX-100) for 15 min at RT, 1X NEBuffer2.1 was added for 1h at 37°C to allow Nt.CviPII-pGL activity to fragment the targeted DNA. For pAG-Mnase activity (1 hr at 4°C or 37°C), buffer including 50 mM Tris.Cl pH 8, 50 mM NaCl and 5 mM CaCl_2_ was used. After 1 wash with 1X PBS, genomic DNA extraction was performed using Monarch genomic DNA purification kit (6 wells pooled per antibody) according to NEB recommendations with a minor modification which included a lysis step at 56°C for 1h instead of 10 min to reverse DNA crosslinking. 0.1 to 1 µg of genomic DNA was used for size selection using first 0.4x volume of NEBNext Sample Purification beads with 10 min incubation at RT. Supernatant containing 200-500 bp DNA sizes was separated from beads using 12-tube magnetic separation rack (NEB # S1509S) and collected. An additional 0.4x volume of NEBNext Sample Purification beads was added to the supernatant and incubated for 10 min at RT. After 2 washes with 80% EtOH, DNA was eluted form beads with 50 µl of low TE buffer and DNA libraries were prepared using NEBNext Ultra II DNA Library Prep (NEB # E7103L) with some modifications. Briefly, end repair/dA tailing was performed on size selected DNA and NEBNext Illumina adaptor (1/10 dilution) were ligated for 2 hrs at RT. USER enzyme (NEB # E7103L) was incubated for 20 min at 37°C and another round of library size selection was performed as described above. The library was amplified using 13 PCR cycles. PCR products were purified using 0.9x volume of NEBNext Sample Purification beads. 1 nM of size selected DNA library was sequenced on Novaseq 6000.

### Immunofluorescence

HT1080 cells were grown on slides (VWR micro cover glass # 48366067) in 6 well plates to 80% confluency. Cells were fixed with 100% MeOH (20 min at -20°C) and incubated for 1h at room temperature with 5% BSA + 1XPBS + 0.3% Triton X-100. Anti-lamin B1 antibody (Abcam # ab8982) was added (1/200 dilution) and incubated for 16 hrs at 4°C. Lamin B1 was then labeled by using an anti-mouse IgG Alexa 488 secondary antibody (Invitrogen # A28175). After 3 washes with 1X PBS + 0.1% Tween 20 for 5 min at RT, anti-lamin B2 (Invitrogen # PA5-29121) was added (1/200 dilution in 5% BSA + PBS + 1X 0.3% Triton X-100) and incubated for 16 hrs at 4°C. Lamin B2 was detected with an anti-rabbit IgG Alexa 594 (Invitrogen # A11012). After 3 washes with 1X PBS + 0.1% Tween 20 (5 min at RT), slides were dried and mounted using Prolong Gold antifade reagent with DAPI (Invitrogen # P36931). Immunofluorescent images were acquired using Zeiss confocal LSM 880 microscope. 3D image of lamin B1 and B2 was constructed using 39 Z-stack images with Zen 2.1 software. Pearson colocalization coefficient of lamin B1 and lamin B2 were determined using Zen 2.1 software. Further intensities for lamin B1/B2 through nuclei diameter were calculated from 40 cells. Average distribution of the intensities was plotted applying Lowess (locally weighted scatterplot smoothing) algorithm using Prism10 (http://www.graphpad.com)

### Hi-C library preparation

HT1080 cells (0.5×10^6^) were crosslinked with 4% formaldehyde for 10 min at RT and quenched with 1.5 M Tris-HCl (pH 8) for 5 min at RT. After 1X PBS wash, cells were permeabilized with Hi-C lysis buffer (10 mM Tris-HCl (pH 8), 10 mM NaCl and 0.2% NP40) for 30 min at 4°C, washed with 1X NEBuffer2 and incubated for 10 min at 58°C with 1X NEBuffer2, 0.3% SDS and 1 mM DTT. To quench SDS, Triton X-100 was added (1% final concentration per well) for 15 min at 37°C. 400 U of MboI (NEB # R0147M) were added and incubated for 16 hrs at 37°C. Enzyme was inactivated at 58°C for 20 min. DNA repair with biotin incorporation as well as ligation were performed accordingly to Liebermann et al. (Liberman et al. 2009, rao et al. 2014). Briefly, DNA polymerase I mix containing NEBuffer2.1 1X (NEB # B7002S), 30 µM of dNTPs (including dCTP/dGTP/dTTP (NEB # N0441S/ N0442S/N0443S respectively) with biotin-14-dATP (ThermoFisher Scientific # 19524016) and 50 U of DNA polymerase I (NEB # M0210M) was added and incubated for 3 hrs at RT. Ligase mix was incubated for 16 hrs at RT. Ligase mix contains 2000 U of T4 DNA ligase (M0202M NEB), 140 µg of recombinant albumin (NEB # B9200S), 1X T4 DNA ligase buffer (NEB # B0202S) and 1 % Triton X-100. After ligation, supernatant was discarded, and cells were washed using 1X PBS. Genomic DNA was extracted from the wells using Monarch Genomic DNA purification Kit (NEB # T3010L). To remove biotin from un-ligated DNA ends, 10 µg of genomic DNA was incubated with 30 U of T4 DNA Polymerase (NEB # M0203L), 25 µM dATP/dGTP (NEB # N0440S/N0442S respectively) in 1X NEBuffer2 for 4 hrs at 20°C. Enzyme was inactivated for 20 min at 75°C. DNA was purified using 1X volume of NEBNext Sample Purification beads (NEB # E6178S) and sheared to 300 bp using Covaris (program : repeat/iterations # 7; repeat process treatment duration 10s; peak power 50W; duty factor 20%; cycles per burst 1000; total treatment time per sample 70s). To capture biotinylated DNA, DNA was incubated with NEB Magnetic Streptavidin beads (NEB # S1420S) for 30 min at RT in 10 mM Tris-HCl (pH 7.5) supplemented with 1 M NaCl. Beads were washed once with 10 mM Tris-HCl (pH 7.5) supplemented with 1 M NaCl and once with low TE buffer and were resuspended in 50 µl of low TE buffer. DNA libraries were prepared on beads using NEBNext Ultra II DNA Library Prep (NEB # E7103L) with some modifications. Briefly, end repair/dA tailing was performed on beads and NEBNext Illumina adaptor (1/10 dilution) were ligated for 2 hrs at RT. 3 µl of USER (NEB # E7103L) was added for 20 min at 37°C. Beads were then washed once with 10 mM Tris-HCl (pH 7.5) supplemented with 1 M of NaCl and once with low TE. Beads were resuspended in 15 µl of low TE. 14 PCR cycles were performed. Supernatant containing PCR products were separated from beads using 12-tube magnetic separation rack (NEB # S1509S) and purified using 0.9x volume of NEBNext Sample Purification beads. 1 nM of DNA Hi-C library was loaded on Novaseq 6000.

### EM-Seq

200 ng of HT1080 genomic DNA was used for detecting methylated cytosine. Protocol was performed according to manufacturer’s recommendations (NEB # E7120S).

### NEED-Seq data processing and peak calling

Sequence analysis was started from Adaptor trimming. Applying Trim Galore, low-quality sequences and adaptor sequences were trimmed (http://www.bioinformatics.babraham.ac.uk/projects/trim_galore) from paired-end sequencing reads using the commend line interface (CLI): --clip_R1 4 –clip_R2 4 –three_prime_clip_R1 4 – three_prime_clip_R2 4. Subsequently, Bowtie2 (Langmead et al., 2012) was applied on trimmed read pairs to map reference genome (hg38) with the following setup: --dovetail –no-unal –no-mixed –no-discordant –very-sensitive -I 0 -X 1000. After getting sam alignment files from Bowtie2, the files were converted to bam files using “samtools view -S -b” (Li et al., 2015) which were further purified by removing PCR duplicates using following settings: “picard MarkDuplicates REMOVE_DUPLICATES=true” and mitochondrial reads (http://broadinstitute.github.io/picard/). Resultant aligned read pairs were utilized for peak calling with MACS2 (Zhang et al., 2008). From NEED-seq, we generated two different set of peaks, (1) Macs2 call peak -f BAMPE -m 4 100 –bdg –SPMR (for narrow peak); (2) Macs2 call peak -f BAMPE -m 4 100 –bdg --broad –SPMR (for broad peak). Finally, FRiP score (fraction of reads in peaks) for each sequence individually were calculated using plotEnrichment (Ramírez et al., 2010).

### Correlation and enrichment analysis

Biological replicates were merged using “samtools merge” (Li et al., 2015). Subsequently, merged bam files were downsized to the same range using Sambamba (Tarasov et al., 2015). Further, the peaks were generated using MASC2 settings as described before (Zhang et al., 2008). Correlation study has two steps 1. Creating .npz file using multiBamSummary 2. Calculate the correlation and plot it as Heatmap or Scatterplot. Deeptools multiBamSummary can create .npz files from a set of bam files and their overall binwise (using bins) distribution or within a specific bed file range (Using BED-file). Here, multiBamSummary was used with BED-file subcommand utilizing the merged peaks of the targeted bam files. Next, deeptools plotCorrelation (--whatToPlot heatmap/ scatterplot) was used for calculating and plotting the correlation coefficient value. Mostly, Pearson and Spearman correlations were applied to calculate coefficient based on requirement. More elaborately, Pearson was used to check similarity based on the co-existence of the reads whereas Spearman represents the quality of the reads within peak regions. Enrichment analysis needs bigwig files. Initially, all the bam files were converted to bigwig files following “bamCoverage –normalizeUsing RPKM” where reads were normalized applying RPKM measure. These bigwig files can further be utilized for signal-to-noise visualization using Integrative Genome Browser (IGV) tools (Robinson et al., 2011). Further, matrices for enrichment were calculated using “deeptools computeMatrix” for both “reference-point” and “scale-regions” modes considering respective peak regions and TSS regions for hg38 downloaded from UCSC genome browser using the generated bigwig files. The resultant matrices were used further as input for “plotProfile” and “plotHeatmap” (Ramírez et al., 2010).

### Other sequencing data study

For comparative study with NEED-seq, data from three established methods were considered: 1. ChIP-seq, 2. CUT&RUN, and 3. CUT&Tag. For ChIP-Seq data, same pipeline was used (already mentioned in NEED-Seq data processing section) with two exceptions. Firstly, subcommand “BAM” was used in the place of “BAMPE” for single end data, and background subtraction was performed while calling the peaks using MACS2 (Zhang et al., 2008). For CUT&RUN and CUT&Tag methods, same NEED-seq pipeline was used.

### Analysis of peaks

Overlapping bed region were defined using “intervene venn” (Khan et al., 2017), demonstrating convergence of bed regions. Bed files were further annotated for their genome-wide localization using annotatePeak.pl from HOMER (Heinz et al., 2010). This tool can predict genomic positions including promoter, intergenic, intron, exon, CpG islands, DNA repeats etc., and it helps to determine the conserved localization for the identified sequences. Also, binding motifs were predicted using the findMotifsGenome.pl from HOMER (Heinz et al. 2010). This tool can either predict the possible binding motifs from different databases e.g., Jasper, GEO etc., and providing the best match (De Novo Motifs) or consider the published information (Known Motifs) as results and corresponding logos.

### Pathway, Gene Ontology Analysis and Bubble plots

Pathways were identified from promoter bound protein coding genes of TFs using EnrichR package of R, considering two criteria 1. p-value ≤ 0.05; 2. At least 10% of the pathway associated genes should belong to the supplied gene list. Bubble plots were made using ggplot2 in R where each bubble represented each pathway and size of the bubble corresponded to -log10(p-value). Here, we have considered top 30 common pathways from TFs where Average(-log10(p-value)) were utilized to determine bubble size.

### Methylation data processing

EM-seq methylation data was used for CpG methylation study. Quality check and removal of the low-quality sequences were performed following the same command which was used in NEED-seq. Further, the fastq files were aligned with reference genome (hg38) using bwameth.py (https://github.com/brentp/bwa-meth). PCR duplicates were removed using “samblaster -e -i” (Faust et al., 2014). Next, mitochondrial reads were removed. Resultant bam files were utilized for further analysis. The methylation call was performed using MethylaDackel extract (github.com/dpryan79/MethylDackel), with coverage 25X threshold and more than 25% per CpG methylation.

### Hi-C Data processing and upstream analysis

For Hi-C data processing, HiC-Pro (V.3.1.0) pipeline was utilized (Servant et al., 2015). HiC-Pro has five processing stages and one quality check stage. For data processing, we have considered MAPQ ≥ 30, MboI ligation sites (GATC) for resolution 50kb and 25kb for hg38 assembly. For ICE normalization, MAX_ITER was provided as 100.

### Identification of LAD domains and HMM states

IgG bam files were utilized for identification of lamin B1 and B2 LAD domains using bamCompare Command Line Interface (CLI): -- scaleFactorsMethod SES –operation log2ratio -bs 20000 (Ramirez et al., 2010, Das et al., 2023). Positive log2 values were considered as LAD domains. Comparative scatter distributions of lamin Bs were performed using bioframe package. Further these lamin B1/B2 LADs were divided in three 25 kb binned states with HMM using Pomegranate (github.com/jmschrei/pomegranate) and Viterbi state calls (github.com/gspracklin/hmm_bigwigs). Predicted states were categorized in three types of domains i.e., strong lamin B1/B2 domains, weak lamin B1/B2 domains, and no LAD domains based on the Jaccard similarity coefficient score between LAD ChIP-seq domains and Lamin Bs states. Highest scores were considered as strong domains, and lowest scores (≥ 0.1) were considered as weak domains and rests were no LAD domains. Next, plotEnrichment was applied to measure read density within strong and weak domains for lamin B1/B2 and genome coverage for all the selected domains were analyzed for NEED-seq lamin Bs and DamID lamin B1 (Meuleman et al., 2013).

### Analysis methodologies for the association of lamin B1/B2 with heterochromatin marks

The association of the heterochromatin marks were identified in two different ways. First, Spearman correlation was calculated for lamin B1/B2 and H3K27me3, H3K9me3, H2AK119ub, H3K9me2 within lamin B domains using MultiBamSummary applying BED-file parameter specification and plotCorrelation for the heatmap (Ramirez et al. 2010). Next, Jaccard similarity between all the histones bound bed regions and lamin B1/B2 strong or weak domains was calculated using “bedtools jaccard” (Quinlan et al., 2010). Furthermore, p-value were calculated to analyze the rate of significance between lamin B1/B2 dependent Jaccard coefficients for each histone marks. In addition, chromHMM based state annotations was performed on lamin B1 and B2 genome-wide domains (Vu et al., 2022).

### Downstream analysis of Hi-C data

HiC-Pro pipeline provided “.matrix” files as well as “.allValidPairs” files as output after ICE normalization (Imakaev et al., 2012). Further “.matrix” was converted in “.mcool” files using hicConvertFormat (Ramirez et al., 2018). “.cool” file with 25kb resolutions were used for further analysis. The “.cool” files were then transformed to obs_exp matrix to study the intrachromosomal interactions using hicTransform and plotted using hicPlotMatrix per chromosome basis (Ramirez et al., 2018). Next, cis interactions and A/B compartments were calculated by applying bioframe and cooltools (Abdennur et al., 2022 (I); Abdennur et al., 2022 (II)). First the .cool files were binned within 25 kb bins and mapped to hg38 chromosome bed regions. Further eigenvalue decomposition-based analysis was performed on “.cool” files to identify cis interactions and followed by Eigenvalue generation for each 25 kb bins using cooler (Abdennur et al., 2020). Positive eigenvalues were considered as A compartments and negative eigenvalues were considered as B compartments.

### Analysis for lamin B1/B2 strong and weak domains within A/B compartments

After calculating the association of lamin Bs with each heterochromatin marks, we have studied lamin B1 and B2 relation with A/B compartment. First Jaccard coefficient was studied between A/B compartments and lamin B1/B2 strong and weak domains. Further overlapping regions of A/B compartments and lamin B1/B2 strong and weak domains were studied for methylation. Per CpG methylation were identified from EM-seq data with ≥ 25x coverage and ≥ 25% per CpG methylation.

### Subcompartments and affinity calculation

Subcompartments were based on Hi-C and NEED-seq lamin B1/B2 data. We confine the Hi-C and lamin Bs with fixed range by using principal component 1 (PC1) from Hi-C (generated using runHiCpca from HOMER; Heinz et al., 2010) and lamin B1 and B2 genome wide domains averaged to 25 kb bins using deeptools MultiBigwigSummary exclusively. Further K-means clustering (K=5) was performed on summarized files separately. The resultant clusters were termed as subcompartments for lamin B1 and B2 (Siegenfeld et al., 2022).

Subcompartments were further utilized to establish the relationship with A/B compartments and histone partitioning. In this case, we have introduced a scoring method called “Affinity score”. Affinity score was calculated based on Z-score of size normalized Jaccard coefficient for each subcompartments and corresponding A/B compartments and heterochromatin marks.

Affinity score (*A*) was defined as:

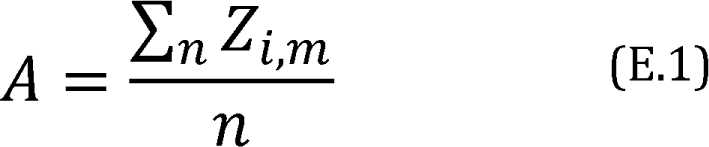

where, *A* is Affinity score between subcompartment *i* and targeted bed regions *m* (where *m* was A/B compartments or Histone marks) for all up-stream and down-stream bed regions padding *n*. Now, *Z_i,m_* is defined as:

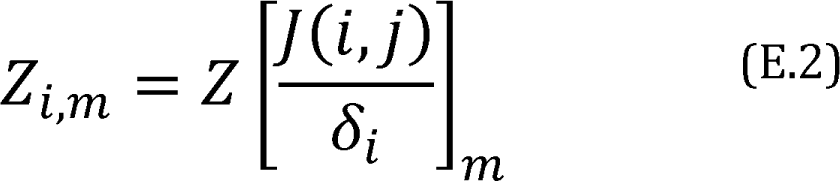

where, *Z_i,m_* is the z-score of size normalized Jaccard value *J* between subcompartment *i* and targeted bed regions *j* for each *m* (where *m* was A/B compartments or Histone marks). The size of each subcompartments was defined as δ_*i*_.

If *m* = histone marks, *n*=4 (no padding, ± 5 kb, ± 10 kb, ± 25 kb),

If *m* = A/B compartment, *n* =1 (no padding)

The final score of *A* can exhibit the affinity of the subcompartments with A/B compartments as well as histone partitioning for each subcompartments. For histone marks, *A* ≥ 0.8 was considered as High affinity partitions and any positive values of *A* < 0.8 were considered as low affinity partitions. For A/B compartments, positive values of *A* were showing rate of affinity of each subcompartments for A/B compartments. Overall, negative *A* values were considered as no affinity.

### Gene ontology analysis

Utilizing the specific histone partitions, we identified promoters closest to the respective subcompartments using “bedtools closest” (Quinlan et al. 2010) where annotation file was downloaded from UCSC genome browser for hg38 assemblies. Further the common and unique promoter bound genes were identified for lamin B1/B2. Next, these selected genes were considered to identify the gene ontology (Biological Processes) using EnrichR library from R interface. Biological processes were selected based on two criteria, 1. p-value ≤ 0.05; 2. At least 3 genes must be associated with the Biological Process.

### RNA-seq data processing

RNAseq data, consisting of three replicates, were analyzed based on STAR aligner protocol. After quality check and adaptor trimming, STAR aligner was used to map the refined fastq with hg38 reference genome (Dobin et al. 2013). Next, all the replicates were merged using samtools. Merged bam were used to identify the transcripts as count matrix applying htseq (Anders et al. 2015) which was normalized based on counts per million (CPM) technique.

### Prediction of polymer structures from Hi-C

Polymer structure was predicted using pHi-C2 package (Shinkai et al., 2022). In this package, the Hi-C interactions modules are converted to polymer network model (a type of Gaussian network model) for a fixed window size. Here we have used 25 kb as window size.

### Data availability

NEED-seq and EM-seq data performed in this study are available in NCBI Gene Expression Omnibus (GEO)-GSE261834.

## Acknowledgments

We thank, T. Evans, D. Comb, Sir R.J. Roberts, J. V. Ellard, and S. Russello for encouragement. The project was funded by basic research grant to SP from New England Biolabs, Inc.

## Author contribution

POE, JB, HGC, GRF, SX performed experiments. VUS and SS performed data analysis. SP conceptualized the project and wrote the manuscript in collaboration with POE, JB, HGC, GRF, VUS, SS and JS.

## Supp. figure legend

**Figure S1.**
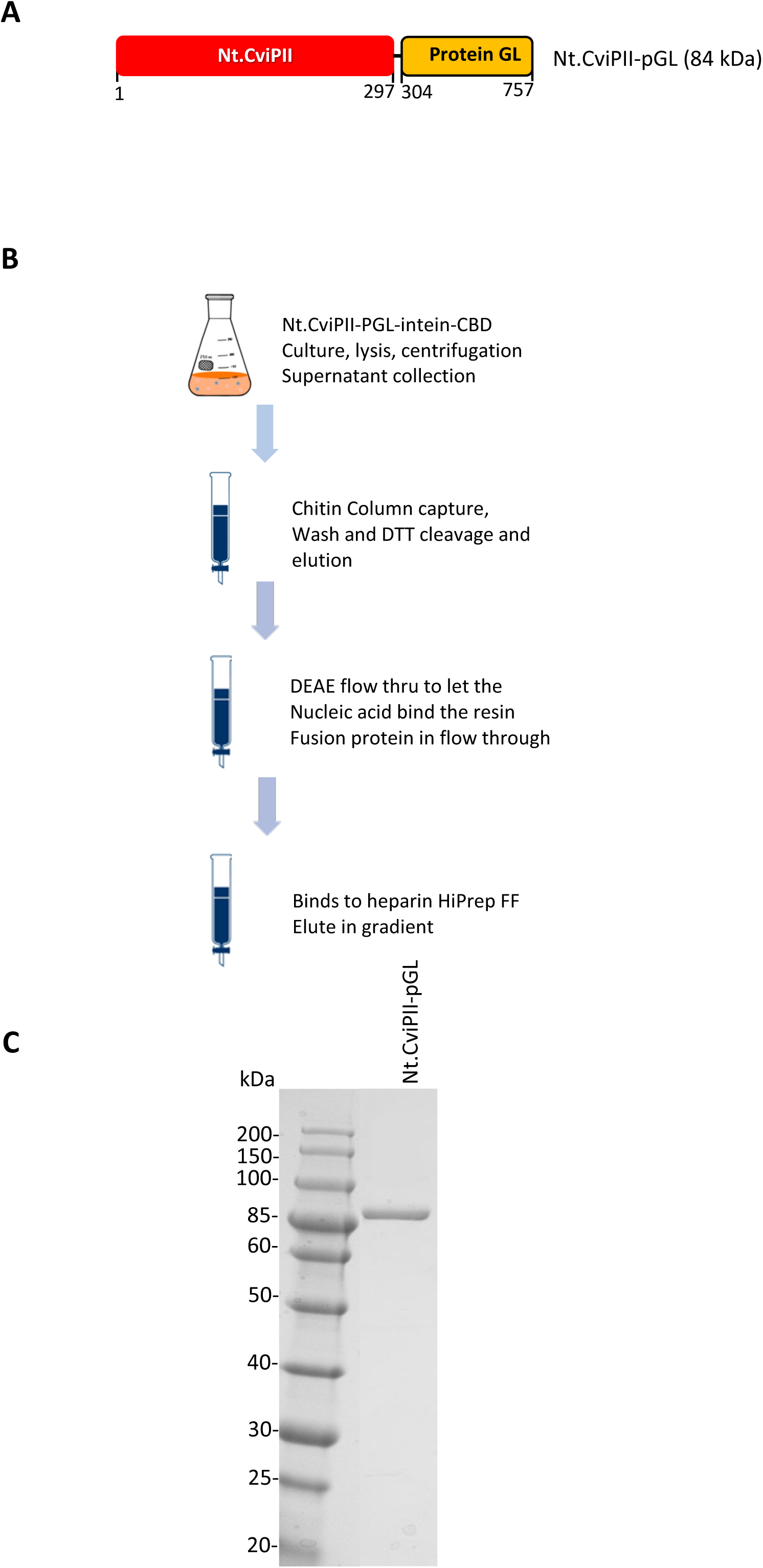
Nt.CviPII-pGL fusion enzyme purification. **(A)** Schematic diagram of Nt.CviPII-pGL fusion enzymes. **(B)** Purification scheme of Nt.CviPII-pGL-intein-CBD fusion. **(C)** Coomassie gel showing Nt.CviPII-pGL purified protein after chitin affinity purification and to homogeneity. The fusion enzyme names are indicated on top.

**Figure S2.**
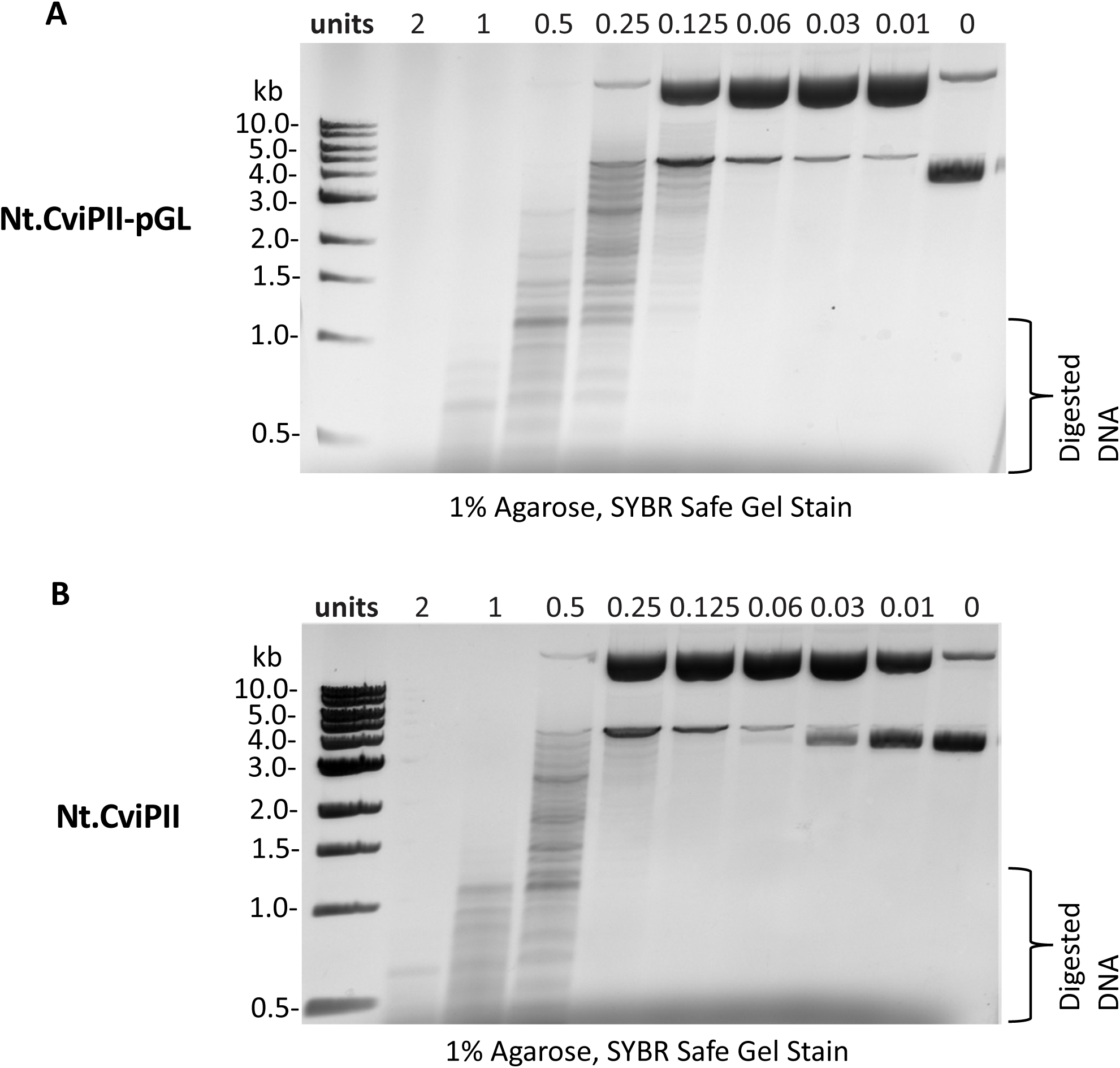
DNA nicking activity assay on pUC19 plasmid. **(A)** DNA nicking activity assay on pUC19 plasmid using different units of Nt.CviPII-pGL fusion enzyme. **(B)** DNA nicking activity assay on pUC19 plasmid using different units of Nt.CviPII (non-fused) enzyme. The DNA fragments were resolved on 1% agarose-TBE gel and visualized under UV.

**Figure S3.**
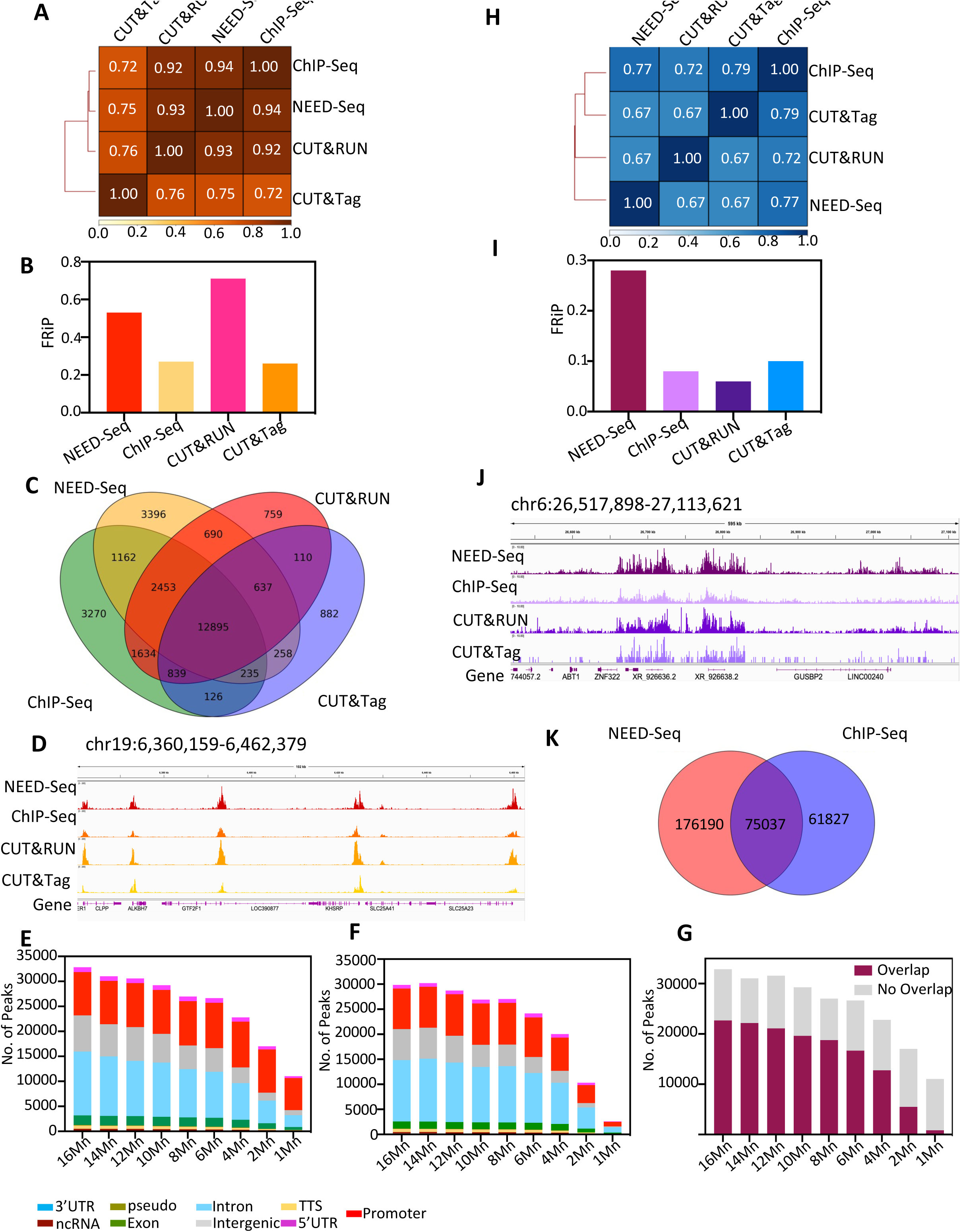
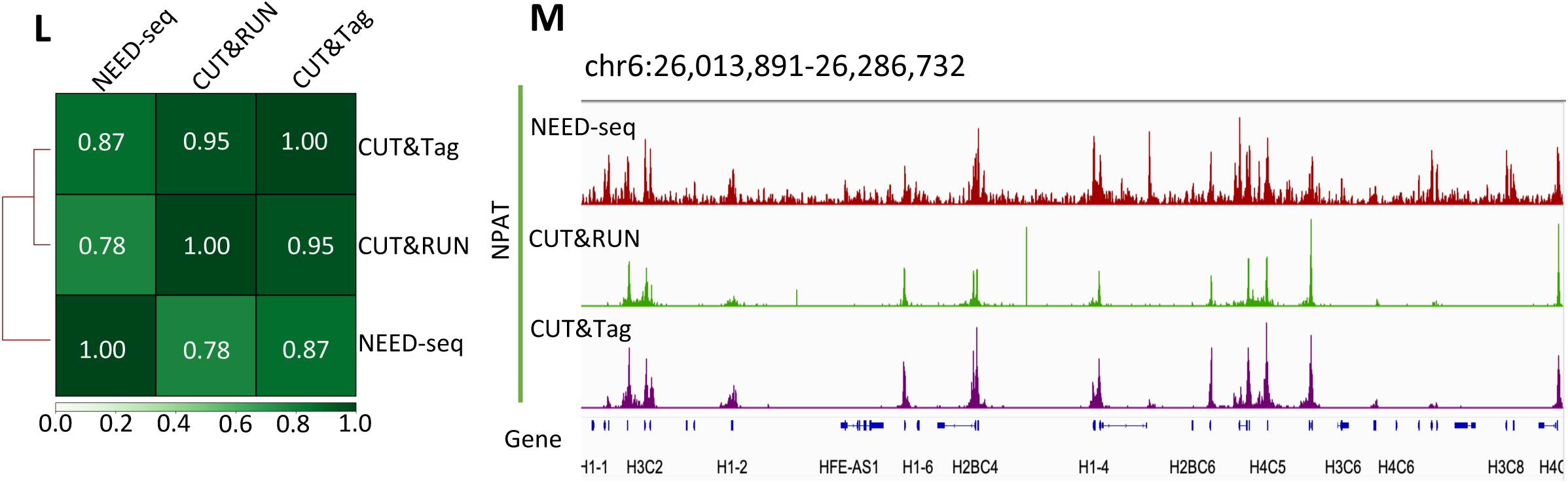
Comparison of NEED-seq, ChIP-seq, CUT&RUN, CUT&TAG of H3K4me3 and H3K27me3 in K562 cells. (**A**) Pearson correlation maps of H3K4me3 NEED-seq with other similar technologies. (**B**) FRiP scores comparison of H3K4me3 NEED-seq with other similar technologies. (**C**) Venn diagram based on common peaks from H3K4me3 sequences from all four methods (**D**) IGV genomic track corresponds to bigwig signals from H3K4me3 (at data range 0-50). (**E**) Feature annotation of H3K4me3 NEED-seq for 16 million (Mn) to 1 Mn reads demonstrating genomic features. (**F**) Similar to E, annotation of H3K4me3 ChIP-seq for 16 Mn to 1 Mn reads demonstrating genomic features as that of E. (**G**) Peak overlaps between ChIP-seq and NEED-seq (at read range 16 Mn to 1 Mn). **(H)** Similar to A, with H3K27me3 NEED-seq Pearson comparison. (**I**) Similar to B, FRiP scores comparison of H3K27me3. (**J**) IGV genomic track corresponds to bigwig signals from H3K27me3 (at data range 0-10). (**K**) Venn diagram based on common peaks from H3K27me3 sequences from ChIP-seq and NEED-seq. Publicly available datasets of CUT & RUN (H4K4me3: GSM3391664 & H3K27me3: GSM2433143), CUT&Tag (H4K4me3: GSM3680226 & H3K27me3: GSM3536511) were used (Table S3). Note CUT&RUN and CUT&TAG was performed using unfixed cells. (**L**) Similar to A and B, with NPAT NEED-seq Pearson correlation comparison between NEED-seq, CUT&RUN, CUT&Tag. (**M**) IGV genomic track corresponding to the bigwig signals from NPAT for NEED-seq, CUT&RUN, CUT&Tag (data range 0-50).

**Figure S4:**
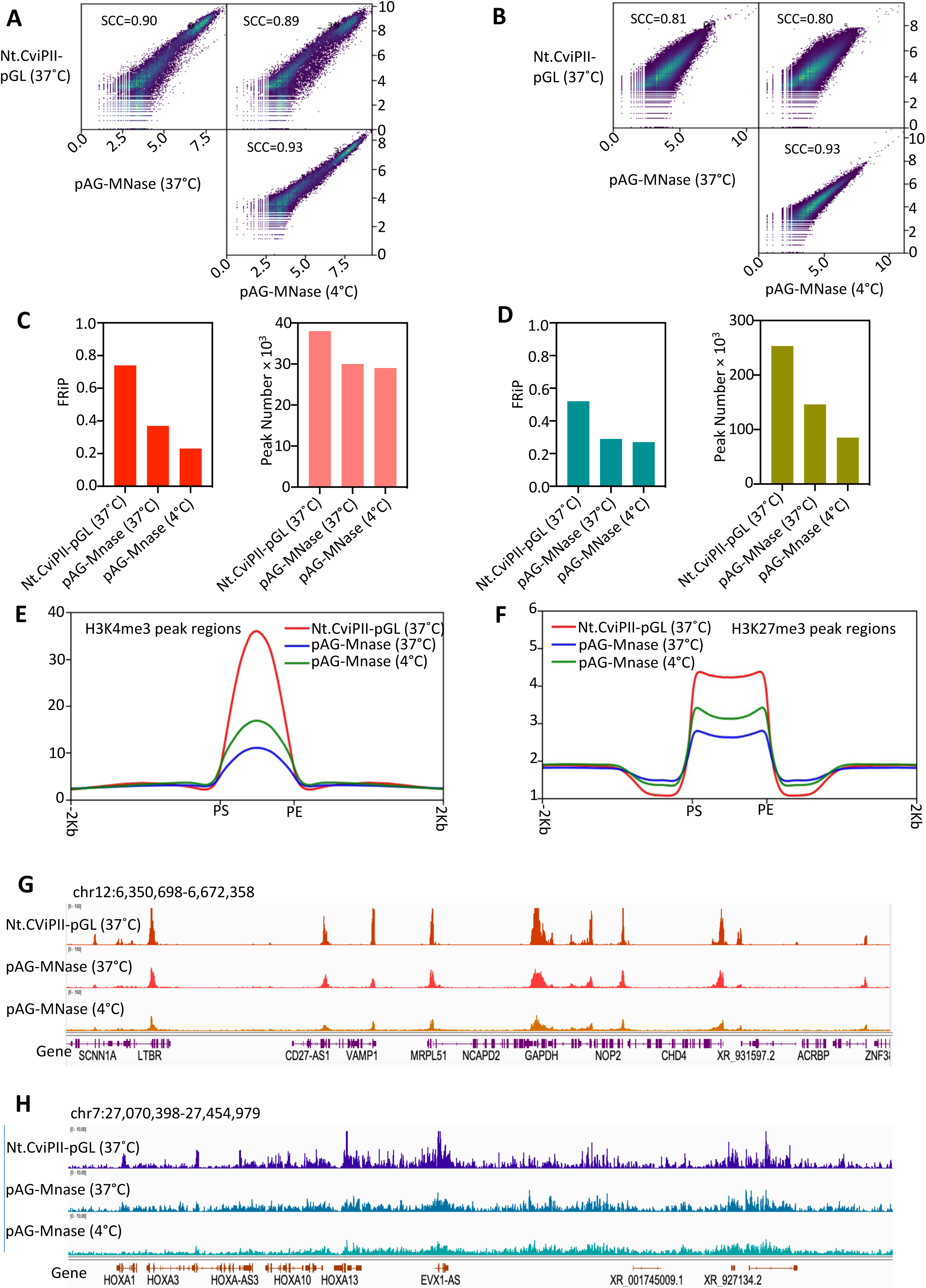
Comparative study between Nt.CviPII-pGL and pAG-MNase in NEED-seq (size selection) application using HT1080 cells. **(A)** Spearman correlation of NEED-seq using anti-H3K4me3 with Nt.CviPII-pGL (37°C) and pAG-MNase (37°C or 4°C). **(B)** Same as A using anti-H3K27me3. **(C)** FRiP-score and peak numbers for H3K4me3 NEED-seq using the enzyme and conditions indicated in X-axis. (**D)** FRiP-score and peak numbers for H3K27me3 NEED-seq using the enzyme and conditions indicated in X-axis. **(E)** Enrichment plots of H3K4me3 peak regions. **(F)** Enrichment plots of H3K27me3 peak regions. **(G)** IGV visualization of H3K4me3 using different enzymes with incubation conditions as indicated. **(H)** IGV visualization of H3K27me3 using different enzymes with incubation conditions as indicated. It may be noted that the recommended condition for pAG-Mnase is 4°C (CUT&RUN) and Nt.CviPII-pGL is 37°C (NEED-seq).

**Figure S5:**
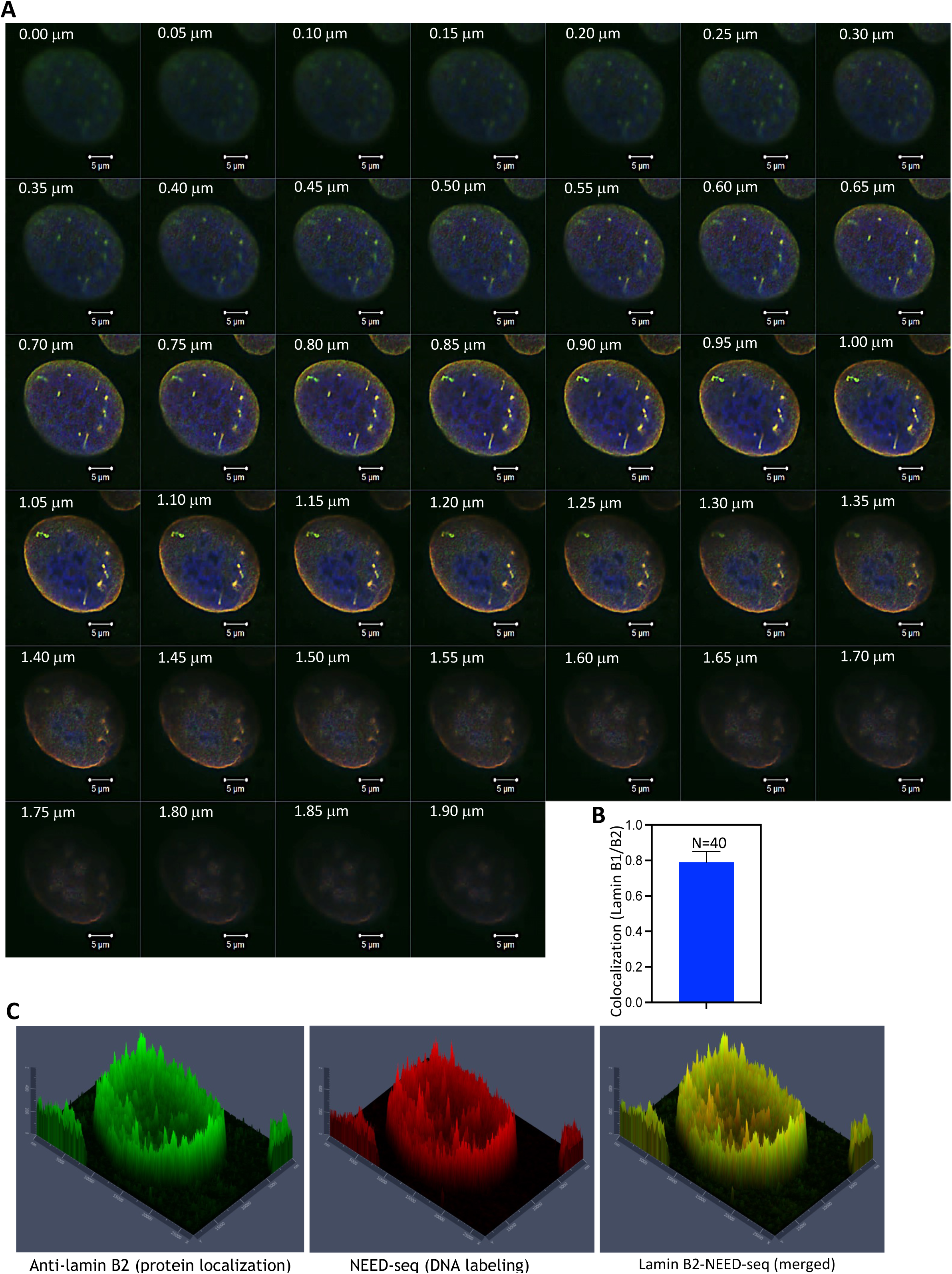
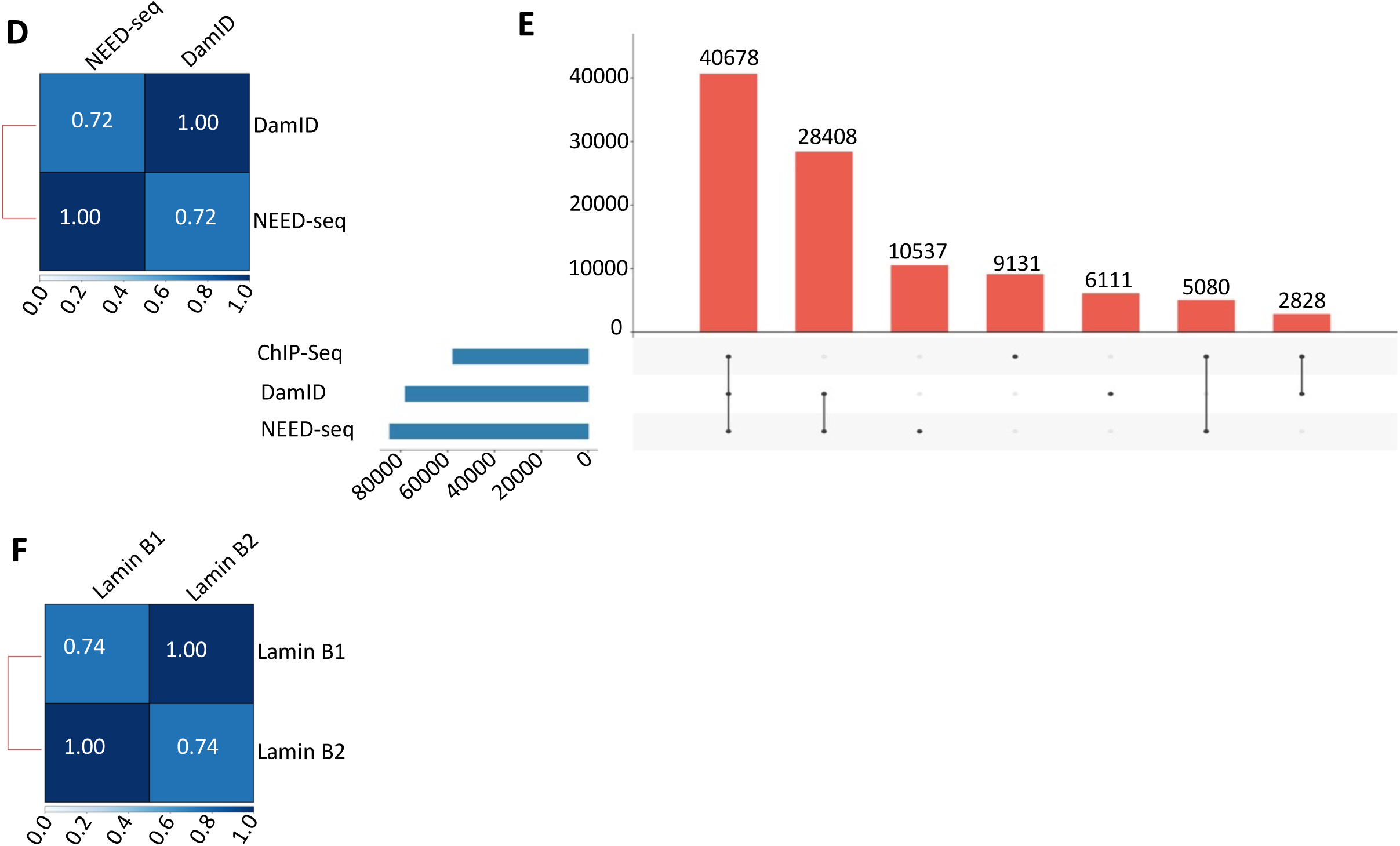
Immunofluorescence of lamin B1 (green) and lamin B2 (red) and lamin B2 NEED-seq in HT1080 cells. **(A)** Image of a cell showing confocal Z-stack images ranging from 0 to 1.9 micrometer. Colocalization is shown in yellow, nucleus stained with DAPI in blue. **(B)** Pearson colocalization coefficient (colocalization) plot of lamin B1 and lamin B2 using 40 cells. (**C**) 3D representation of microscopic images exhibiting anti-lamin B2 labelling (green), NEED-seq DNA labelling (red), and merged (yellow) representation. **(D)** Spearman Correlation between lamin B1 NEED-seq and DamID (r=0.72). **(E)** UpSet plot between lamin B1 bound domains (20 kb) from ChIP-seq, DamID and NEED-seq demonstrating common regions. **(F)** Spearman correlation between lamin B1 and lamin B2 (r=0.74).

**Figure S6:**
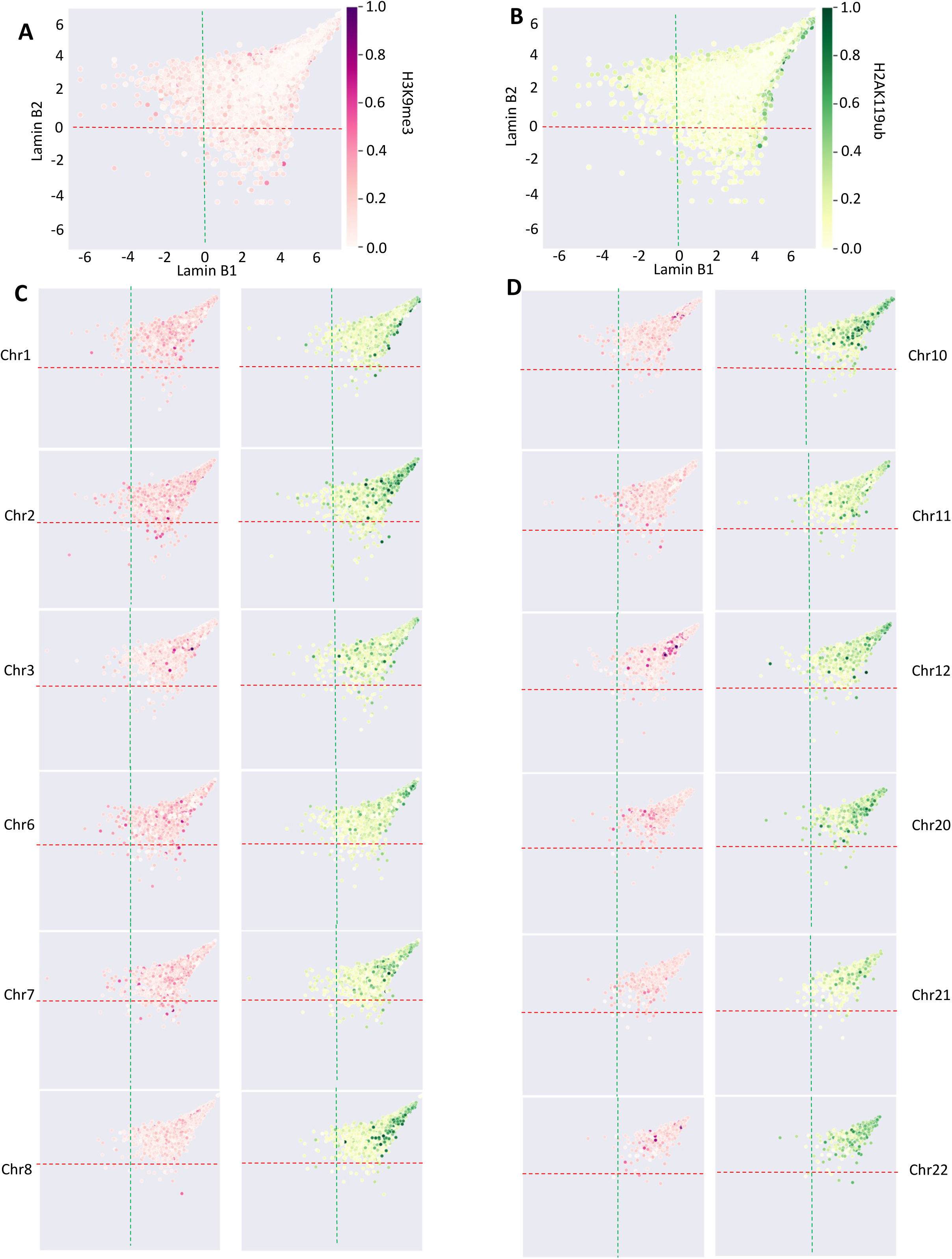
Visualization of lamin B1/B2 domain distribution with H3K9me3 and H2AK119ub signal intensity derived from NEED-seq. **(A)** Scatter plot showing genome-wide distribution of H3K9me3 bound to lamin B2 in red **(B)** Scatter plot showing genome-wide distribution of H2AK119ub bound to lamin B1 in green **(C-D)** Chromosome-wise distribution for chr1-3, chr6-8 and chr10-12, chr20-22 were shown for H3K9me3 (red) and for H2AK119ub (green).

**Figure S7:**
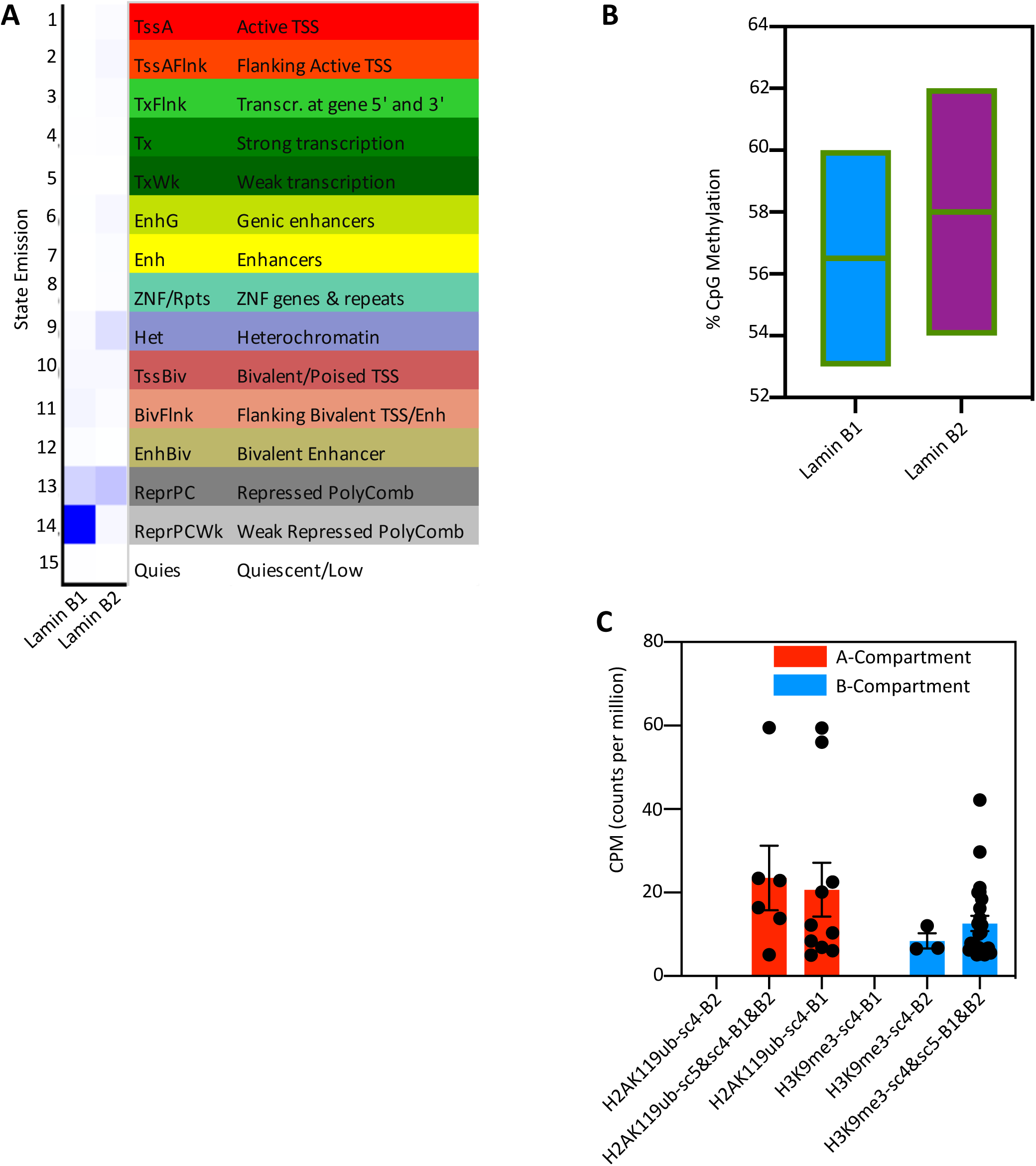
Lamin B1/B2 state emission and DNA methylation. **(A)** ChromHMM states for lamin B1/B2 domains **(B)** Distribution of CpG methylation at B-compartment bound lamin B1 and lamin B2. Both strong and weak domains were used in this analysis. (C) Genes with H2AK119ub or H3K9me3 marks from sc4 and 5 mapped (based on fig. 6G and H) to A and B compartment respectively, correlated to RNA expression profile in CPM.

**Figure S8:**
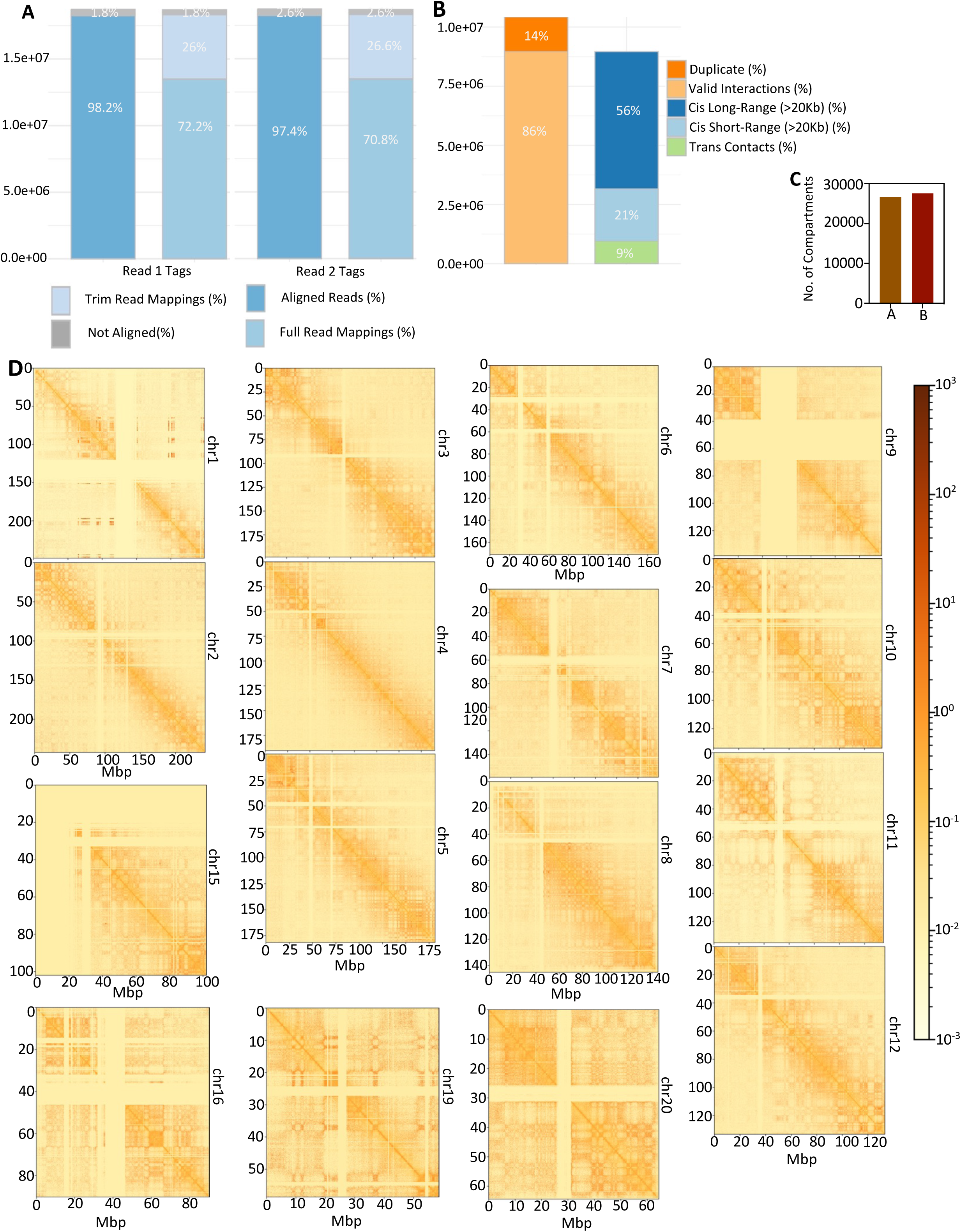
Data analysis and quality matrix of Hi-C of HT1080 using hg38. **(A)** QC analysis of Read specific mappability. **(B)** Statistical study on interaction pairs. **(C)** Number of domains within A/B compartments for 25 kb resolution. **(D)** Chromosome-wise interaction maps based on obs_exp matrix.

**Table S1:** List of histone marks from HT1080 cell line and corresponding FRiP score obtained.

**Table S2:** List of Pathways associated with (**A**) AP1 promoter positive genes; (**B**) RB1_p53 promoter positive genes.

**Table S3:** List of peaks identified with H3K27me3 from NEED-seq, ChIP-seq, CUT&RUN and CUT&Tag.

**Table S4:** List of Biological processes associated with promoter bound lamin B1/B2 domains **(A)** For H2AK119ub subcompartments **(B)** For H3K9me3 subcompartments.

**Table S5:** List of antibodies used in NEED-seq.

**Table S6:** Details on publicly available data utilized for analysis.

