## Supplementary material for "Distinct structural and functional heterochromatin partitioning of lamin B1 and B2 revealed using genome-wide Nicking Enzyme Epitope targeted DNA sequencing": Table S1

| Serial No. | Histone Marks | FRiP Score |
| --- | --- | --- |
| 1 | **H3K9me3** | **0.202** |
| 2 | **H2AK119ub** | **0.375** |
| 3 | **H4K20me3** | **0.243** |
| 4 | **H3K27me3** | **0.25** |
| 5 | **H3K27ac** | **0.323** |
| 6 | **H3K4me3** | **0.326** |
| 7 | **H3K4me1** | **0.377** |
| 8 | **H4K20me1** | **0.273** |

**Table S1**: List of Histone marks from HT1080 cell line and corresponding FRiP score
