## Supplementary material for "Distinct structural and functional heterochromatin partitioning of lamin B1 and B2 revealed using genome-wide Nicking Enzyme Epitope targeted DNA sequencing": Table S3

**Table S3: Comparison of number of Peaks for H3K27me3 for NEED-seq and other methods**

| **Method** | **Peak Number** |
| --- | --- |
| NEED-Seq | 251,227 |
| ChIP-Seq | 276,405 |
| CUT and RUN | 19,124 |
| CUT and TAG | 17,452 |
