## Supplementary material for "Distinct structural and functional heterochromatin partitioning of lamin B1 and B2 revealed using genome-wide Nicking Enzyme Epitope targeted DNA sequencing": Table S6

| **Cell line** | **Samples** | **Method** | **Read Alignment Type (PE/SE)** | **Assembly** | **Source** | **Accession** |
| --- | --- | --- | --- | --- | --- | --- |
| K562 | H3K4me3 | CUT&RUN | PE | hg38 |  | GSM3391664 |
| K562 | H3K4me3 | CUT&Tag | PE | hg38 |  | GSM3680226 |
| K562 | H3K27me3 | CUT&RUN | PE | hg38 |  | GSM2433143 |
| K562 | H3K27me3 | CUT&Tag | PE | hg38 |  | GSM3536511 |
| K562 | DNAse-Seq | DNAse-Seq | PE | hg38 | ENCODE Project Consortium | [GSM2400371](https://www.ncbi.nlm.nih.gov/geo/query/acc.cgi?acc=GSM2400371) |
| HT1080 | Open Chromatin | NiCE-Seq | PE | hg38 |  | GSM4285582, GSM4285583 |
| K562 | NPAT | CUT&RUN | PE | hg38 |  | GSM3560265 |
| K562 | NPAT | CUT&Tag | PE | hg38 |  | GSM3609774 |
| HT1080 | Lamin B1 | DamID | SE | hg38 |  | GSE87148 |

Table S6: List of publicly available data utilized in the analysis
